# Multi-parameter spectral flow cytometry panel for immune phenotyping of murine B and T cell responses

**DOI:** 10.1101/2025.07.18.665522

**Authors:** Kassandra Hoetzel, Hendrik Feuerstein, Julia Ludwig, Hedda Wardemann

## Abstract

To enable the analysis of B cell and T cell immune responses in mouse lymph nodes, spleen, and bone marrow, including different B cell and T cell subsets and their activation status, as well as the antigen-reactivity and isotype of B cells and level of T cell exhaustion, a novel 31-parameter spectral flow cytometry panel was developed.

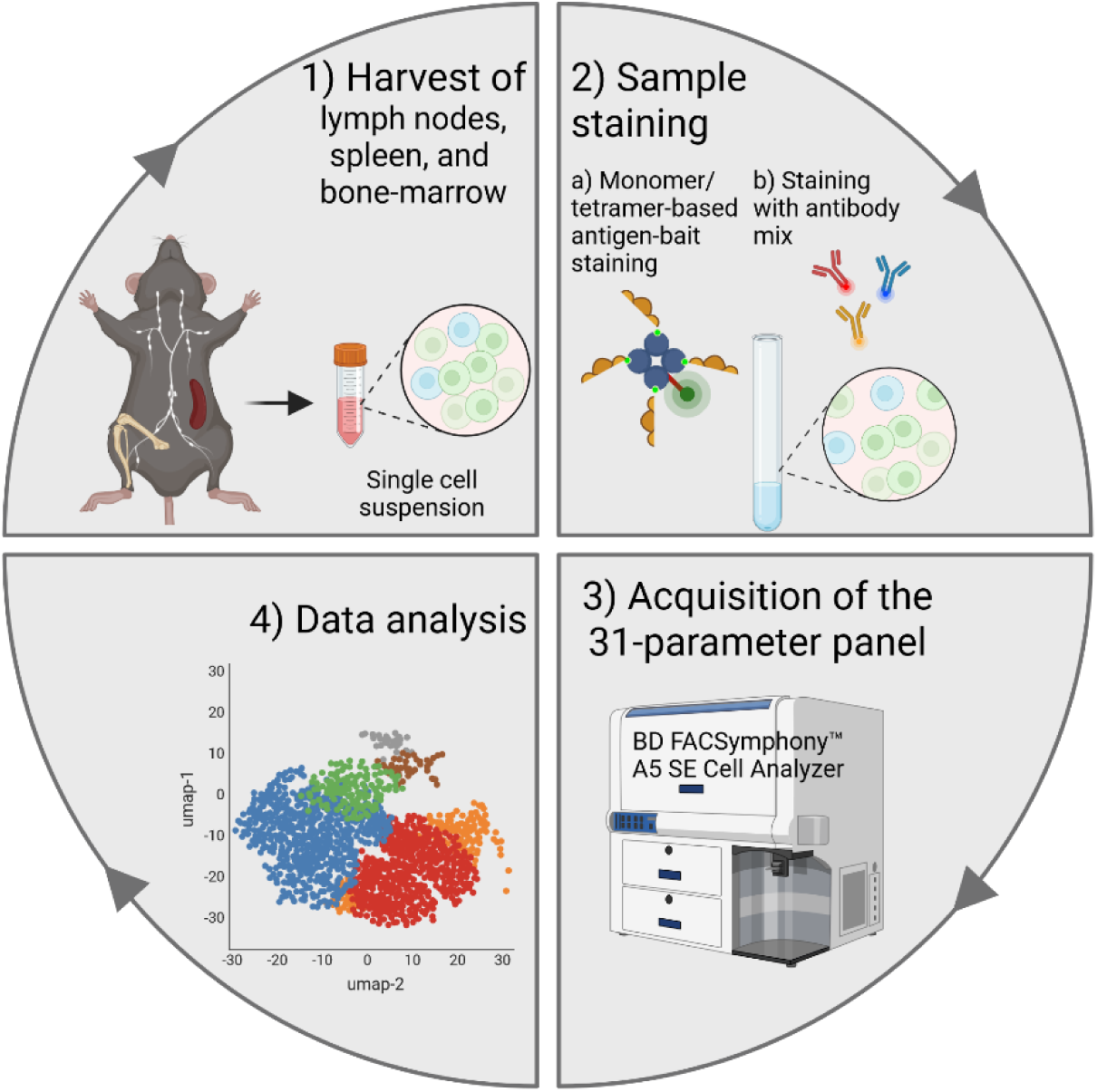

Created in https://BioRender.com

---

The primary site of T cell dependent B cell immune responses are germinal centers in secondary lymphoid organs [1]. Within these microanatomical structures, antigen-activated B cells undergo affinity maturation, a process that selects somatically mutated B cells with improved antigen-binding based on the competition for antigen and T cell help driving differentiation into antibody-secreting and quiescent memory B cells [1]. [2]. During recall responses, antigen-binding induces the rapid differentiation of memory B cells into antibody-secreting cells contributing to the fast production of antigen-specific serum antibodies. The majority of newly developing antibody-secreting cells that develop during primary and secondary immune responses are short-lived plasmablasts but a fraction can occupy dedicated survival niches in bone marrow differentiating into long-lived plasma cells mediating durable humoral immunity.

Studies in mice continue to provide important insights in the cellular and molecular mechanisms underlying B cell responses. However, deep cellular analyses are often limited by the low number of antigen-specific cells that participate in immune responses in individual animals [2]. High-dimensional flow-cytometric analyses enable the parallel analysis of diverse cell populations if cell numbers are a limiting factor, for example when analyzing antigen-specific immune responses in murine lymph node samples [3–14]. We therefore established a multicolor spectral flow cytometry panel using a full visible spectrum cytometer (BD FACSymphony™ A5 SE Cell Analyzer) equipped with five lasers and 49 detectors, facilitating the deep parallel phenotypic assessment of B cells and T cells in lymph nodes, spleen, and bone marrow with a specific focus on antigen-experienced subsets (Tables S1 and S2, Fig. S1).

Here, we report the development and validation of this 31-parameter panel using lymph node, spleen and bone marrow samples from non-immunized mice and mice immunized with trimeric SARS-CoV-2 prefusion spike protein stabilized by six prolines (S6P). Antigen-reactive B cells were identified using fluorochrome-labeled SP6 and SARS-CoV-2 receptor-binding domain (RBD) baits. The materials and methods, including the antibody list (Table S3), single color reference controls (Table S4), the antibody titration process (Fig. S2), and the panel design and optimization (Table S5 and S6) are described in the Supplementary Information.

Independent of the origin of the sample, single lymphocytes in all organs were identified based on their size and granularity (Fig. 1A and 2A). Dead cells as well as NK cells, erythrocytes, monocytes including neutrophils, macrophages, and dendritic cells (collectively defined as lineage negative; Lin-) were excluded from the analysis using the fixable viability stain 440UV and anti-NK-1.1, anti-TER-119, anti-Ly-6G/Ly-6C (Gr-1), anti-F4/80, and anti-CD11c antibodies all labeled with the same fluorochrome, respectively (Fig. 1A and 2A) [3]. Among single live lymphocytes, B cells were identified based on the lineage markers CD19 and B220. In the spleen antibodies against CD93, CD21, and CD23, enabled the discrimination of transitional B cell subsets (T1 – CD93^high^IgD^neg/low^CD23^-^; T2/T3 – CD93^low^IgD^low/pos^CD23^+^) from marginal zone (IgM^+^CD21^+^CD23^low^) and mature naïve (CD23^high^CD21^+^) B cells (Fig. 1B, C) [4, 5, 8]. In immunized but not in non-immunized mice, a small fraction of mature naïve cells expressed the activation marker CD80 but not CD86 (Figure 1D and Fig. S4).

**Figure 1:**
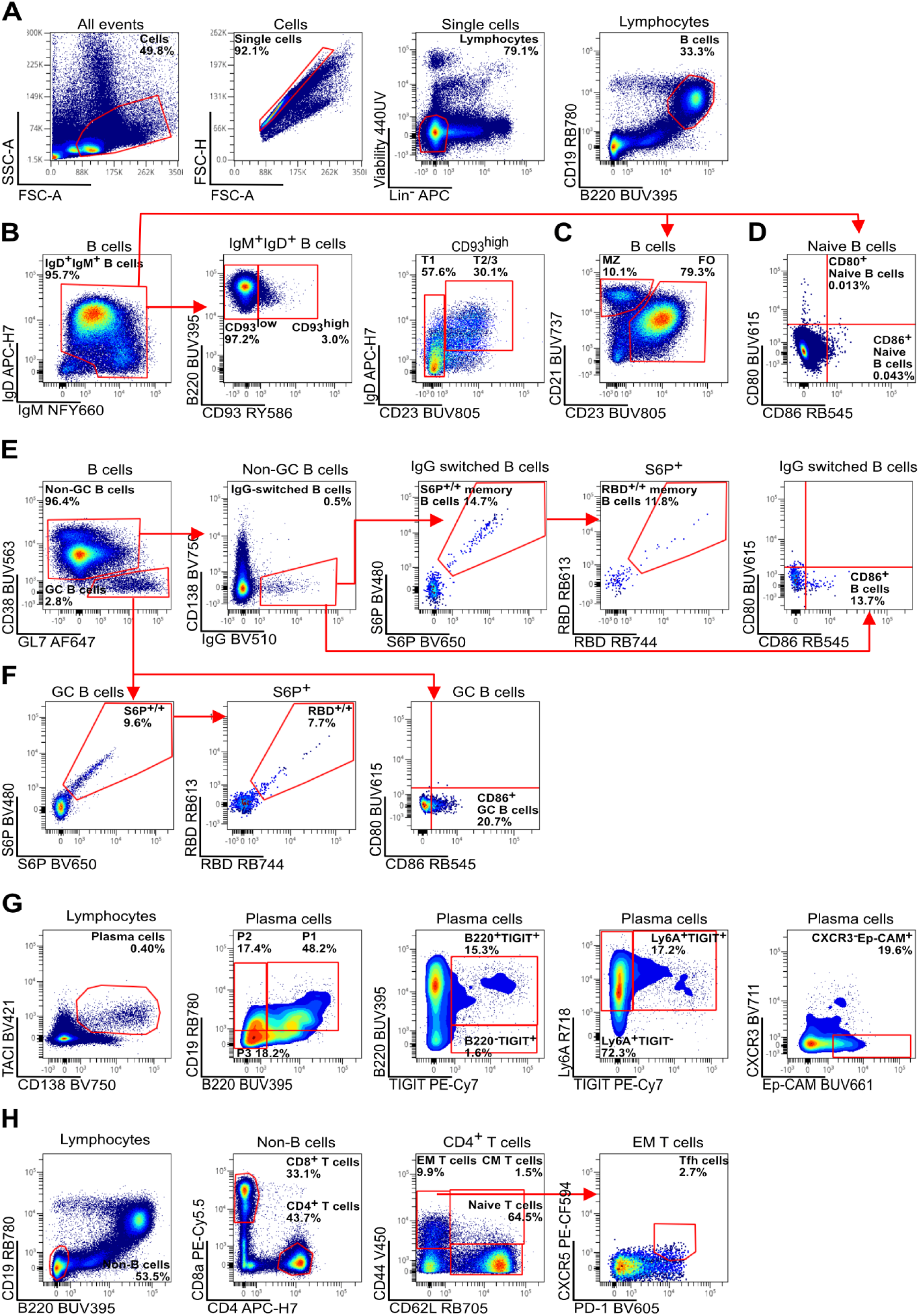
Gating strategy for murine spleen samples. A single cell suspension was generated from the spleen of an immunized mouse 7 days after secondary immunization. **(A)** Gating strategy for viable lymphocytes after exclusion of cellular debris, doublets, dead cells, and erythrocytes (TER^-^119^+^), monocytes and neutrophils (Ly-6G/Ly-6C^+^ (Gr-1^+^)) as well as macrophages (F4/80^+^), dendritic cells (CD11c^+^), and NK cells (NK-1.1^+^). B cells (CD19^+^B220^+^) were identified based on the surface expression of CD19 and B220. **(B)** From IgM^+^IgD^+^ B cells, CD93 (AA4.1)^high^ B cells were further gated to identify T1 stage transitional B cells (IgM^high^CD23^low^) and T2 stage transitional B cells (IgD^high^CD23^high^). **(C)** Mature naive and marginal zone B cells were defined as CD23^high^CD21^+^ and CD23^low^CD21^high^, respectively. **(D)** IgM+IgD+ B cells were analyzed for expression of activation markers CD80 and CD86. **(E)** To identify memory B cells, plasma cells (PCs; CD138^+^) were excluded from non-GC B cells and further gated on class-switched B cells (IgG^+^). Among these IgG^+^ class-switched B cells, antigen-reactive (S6P^+/+^, RBD^+/+^) cells were identified as well as activation marker (CD86^+^) expressing class-switched B cells. **(F)** GC B cells (CD38-GL7+) were identified based on the lack of CD38 expression. GC B cells were investigated for antigen-reactivity (S6P^+/+^, RBD^+/+^) as well as their activation status (CD86^+^). **(G)** PCs and plasmablasts (PBs; TACI^int^CD138+) were divided into subsets P1 – early dividing precursor PCs (B220^int^CD19^int^), P2 - early PCs (B220^low^CD19^int^), and P3 - mature resting PCs (B220^low^CD19^low^). Expression of TIGIT (and B220) was indicative of early and likely still proliferation P1 and P2 antibody secreting cells. CXCR3, Ep-CAM and Ly6A, markers were associated with IgG (Ep-CAM^high^CXCR3^-^) or IgA (Ly6A^high^TIGIT^-^) expression. **(H)** Among non-B cells, CD8^+^ and CD4^+^T cells were discriminated. CD4^+^ T cells were separated into EM T cells (CD44^+^CD62L^-^), CM T cells (CD44^+^CD62L^+^), and naive T cells (CD44^-^CD62L^+^). EM T cells were further gated to identify Tfh cells (CXCR5^+^PD-1^+^).

**Figure 2:**
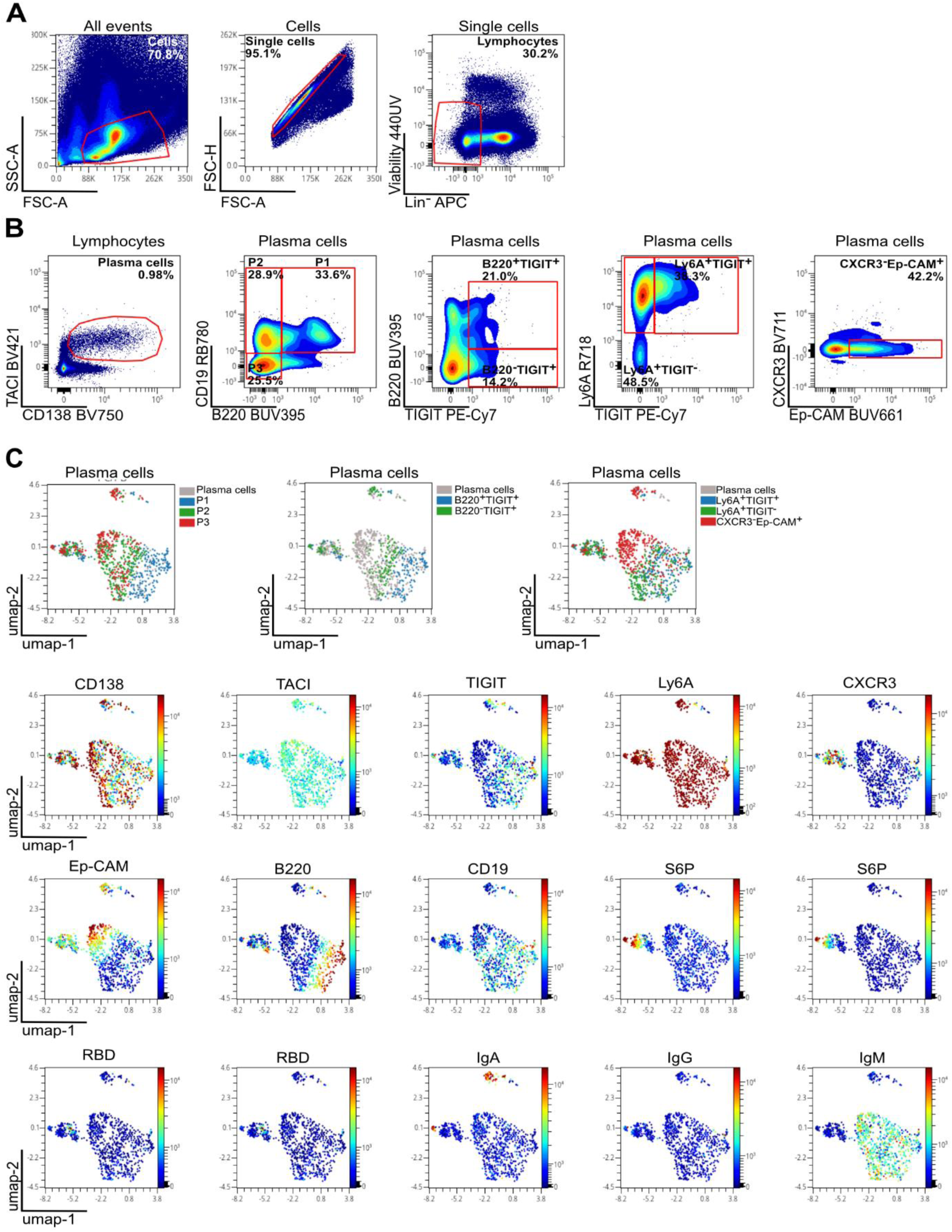
Gating strategy for murine bone-marrow plasma cell subsets. Single cell suspension from bone marrow of an immunized mouse 7 days after secondary immunization. **(A)** Gating strategy for viable lymphocytes after exclusion of cellular debris, doublets, dead cells, and erythrocytes (TER^-^119^+^), monocytes and neutrophils (Ly-6G/Ly-6C^+^ (Gr-1^+^)) as well as macrophages (F4/80^+^), dendritic cells (CD11c^+^), and NK cells (NK-1.1^+^). **(B)** PCs and PBs (TACI^int^CD138+) were divided into subsets P1 – early dividing precursor PCs (B220^int^CD19^int^), P2 - early PCs (B220^low^CD19^int^), and P3 - mature resting PCs (B220^low^CD19^low^). Expression of TIGIT (and B220) was indicative of early and likely still proliferation P1 and P2 antibody secreting cells. CXCR3, Ep-CAM and Ly6A, markers were associated with IgG (Ep-CAM^high^CXCR3^-^) or IgA (Ly6A^high^TIGIT^-^) expression. **(C)** UMAP analysis of the final 31-parameter flow cytometry panel. For the dimensionality reduction approach, cellular debris, doublets, dead cells, and erythrocytes (TER-119+), monocytes and neutrophils (Ly-6G/Ly-6C+ (Gr-1+)) as well as macrophages (F4/80+), dendritic cells (CD11c+), and NK cells (NK-1.1+) were excluded. UMAP was applied on the PC gate using OMIQ (Nearest Neighbors = 50, Minimum Distance = 0.1, Metric = Euclidean, Components = 2).

CD38 was expressed on marginal zone, mature naïve, and IgG^+^ class-switched B cells in the spleen (Fig. 1E, Fig. S5). To better separate the IgG^+^ class-switched B cell population and also exclude plasma cells, antibodies against CD138 and IgG were used (Fig. S6 and S7). The frequency of IgG^+^ class-switched B cells was higher in immunized mice including IgG^+^ cells that bound antigen (S6P, RBD) and expressed the activation marker CD86 reminiscent of memory B cells (Fig. 1E and S4). GL7 expressing CD38-negative GC B cells that were rare in non-immunized mice but clearly detectable in spleen and lymph nodes of immunized mice (Figure 1F and Figs. S4, S8, S9) included a sizable fraction of antigen (S6P, RBD)-reactive cells, as well as cells with high expression of CD86, reminiscent of GC light-zone B cells (Figure 1F) [1, 7–10]. Immunization also led to the development of plasma cells, which were identified by the expression of CD138 and the transmembrane activator and CAML interactor (TACI) independent of their levels of B220 and CD19 that are downregulated during plasma cell maturation (Fig. 1G and 2A-B, Fig. S4 and S10). [11–13]. In spleen and bone marrow, three plasma cell subsets (P1-3) were discriminated based on their CD19 and B220 expression: P1 - plasma cell precursors (B220^int^CD19^int^), P2 - early plasma cells (B220^lo^CD19^int^), and P3 - mature plasma cells (B220^low^CD19^low^). Our panel includes antibodies against TIGIT, expressed predominantly on early and likely still proliferation P1 and P2 antibody secreting cells, as well as CXCR3, Ep-CAM and Ly6A, markers associated with IgG (Ep-CAM^high^CXCR3^-^) or IgA (Ly6A^high^TIGIT^-^) expression [12, 13].

To facilitate the assessment of T cell responses, antibodies against CD4, CD8, and CXCR5 - expressed on T follicular helper (Tfh) cells - as well as the activation marker PD1 were included in the panel (Figure 1H) [1, 8, 14]. Based on the expression of CD44 and CD62L, naïve (CD44+CD62L+), effector memory (EM; CD44+CD62L-) and central memory (CM; CD44-CD62L+) T cells were distinguished. Tfh cells were rare and expression of CXCR5 was overall low compared to B cells, even after immunization (Fig. S11 and S12).

Uniform Manifold Approximation and Projection (UMAP) is a nonlinear dimensionality reduction technique that enables the visualization of complex high-dimensional data in two dimensions while preserving the global structure of the dataset. UMAP accelerated the complex and time-intensive analysis of the high-dimensional datasets generated with our panel (Fig. 2C, Fig. S11 and S12). The analysis confirmed and helped to visualize the identity of the different B cell and T cell subsets.

In summary, our 31-parameter spectral flow cytometry panel allows the phenotypic discrimination of diverse B cell and T cell lineages with a special focus on B cell subsets in immunized mouse lymphoid tissue and bone marrow including antigen-reactive B cells.

## Abbreviations

CM: Central memory
EM: Effector memory
FO: Follicular
GC: Germinal center
MZ: Marginal zone
PB: Plasmablast
PC: Plasma cell
RBD: SARS-CoV-2 receptor-binding domain
S6P: SARS-CoV-2 prefusion spike protein stabilized by six prolines
TACI: Transmembrane activator and CAML interactor
Tfh: T follicular helper (cell)
UMAP: Uniform Manifold Approximation and Projection

## Acknowledgements

The authors thank Steffen Schmitt (Core Facility for Flow Cytometry, DKFZ) and Sandra Blaszkiewicz (Field Application Support Manager, BD Life Sciences - Biosciences) for technical support during panel optimization. We thank the Vaccine Formulation Institute (VFI, Geneva, Switzerland) for providing both the SARS-CoV-2 prefusion spike protein stabilized by six prolines and the liposome-based adjuvant (LMQ). Hedda Wardemann designed the study, interpreted data, and wrote the manuscript. Kassandra Hoetzel designed the study, performed experiments, interpreted data, and wrote the manuscript. Hendrik Feuerstein and Julia Ludwig performed experiments and interpreted data.

## Funding information

This work was supported by the Helmholtz Association’s Initiative and Networking Fund project “Virological and immunological determinants of COVID-19 pathogenesis—lessons to get prepared for future pandemics (KA1-Co-02_CoViPa).” Hendrik Feuerstein and Kassandra Hoetzel were supported by the DKFZ International PhD Program.

## Conflict of Interest

The authors declare no commercial or financial conflict of interest.

## Open Research

### Data availability statement

The flow cytometry files are available from the corresponding author upon reasonable request.

### Ethics approval statement

All C57BL/6J mouse procedures were approved by the Regierungspräsidium Karlsruhe, Germany (project number G27/23) and experiments were conducted in accordance with the German Animal Protection Law.

## Supplementary Information Supplementary Materials and Methods

Description of the materials and methods used to prepare single cell suspensions from lymph nodes, spleen, and bone marrow.

### Material

#### Commercially available reagents

- PBS (Gibco, Cat. 70011-036)
- Fetal bovine serum (FBS) (Sigma-Aldrich, Cat. TMS-013)
- Dimethylsulfoxid (Sigma-Aldrich, Cat. D2650)
- 40 µM nylon cell strainer (Corning, Cat. 431750)
- Cryogenic vial (Corning, Cat. 431417)
- Corning® CoolCell™ FTS30 (Corning, Cat. 432006)
- BD Brilliant Buffer (BD Biosciences, Cat. 566349)
- RPMI 1640 (Gibco, Cat. 21875034)
- Dulbecco’s Phosphate Buffered Saline (DPBS) (Gibco, Cat. 14190-144)
- Neubauer counting chamber (Millipore, Cat. MDH-2N1-50PK)
- Trypan blue stain 0.4% (Life Technologies, Cat. T10282)
- Fixable Viability Stain 440UV (BD Biosciences, Cat. 566332)
- Fc-Block – purified anti-mouse CD16/32 antibody (Invitrogen, Cat. 14-0161-82)
- CellBlox™ blocking buffer (Thermo Fisher Scientific, Cat. B001T06F01)

#### Buffers

<u>Freezing medium</u>: 80% (v/v) Fetal bovine serum, 20% (v/v) Dimethylsulfoxid <u>Staining buffer</u>: 2% (v/v) Fetal bovine serum in PBS

<u>Live-Dead dye solution</u>: Dilute fixable Viability Stain 440UV 1:500 in Dulbecco’s Phosphate Buffered Saline

Brilliant staining buffer: Mix 1 part of BD Brilliant Buffer with 1.8 parts of Staining buffer

### Methods

*Mice (C57BL/6J, aged 6–8 weeks) received two intramuscular immunizations, four weeks apart, with SARS-CoV-2 prefusion spike protein stabilized by six prolines (S6P) formulated in liposome-based adjuvant (LMQ); or were left non-immunized as controls (Hoetzel et al., unpublished). Mice were sacrificed one week after the second immunization, and samples were processed and cryopreserved as described below*.

*Lymph node and spleen processing protocol:*

1. Dissected inguinal, axial, and popliteal lymph nodes or spleen (for titration and cell compensation) from immunized mice and transferred them into falcons with PBS.
2. Pressed tissue pieces through a 40 µm cell strainer into the same falcon filled with PBS using a syringe plunger.
3. Centrifuged at 403 × g for 10 minutes at 4°C.
4. Removed the supernatant, resuspended lymph node cells in 400 µl FBS, counted cells, and added 400 µl freezing medium (3 million cells per tube). Removed the supernatant, resuspended splenocytes in 900 µl FBS, counted cells, and added 900 µl freezing medium (6 million cells per tube).
5. Froze cells immediately in a Corning® CoolCell™ FTS30 at -80°C overnight.
6. Transferred cells to liquid nitrogen the next day.

*Bone marrow processing protocol:*

1. Prepared tibia and femur from immunized mice with scissors and cut the bones open at the ends.
2. Placed the bones into an Eppendorf tube and centrifuged at 956 × g for 6 minutes at 4°C to flush out the cells.
3. Removed the bones and resuspended the pellet in 500 µl FBS.
4. Centrifuged again at 956 × g for 6 minutes at 4°C and removed the supernatant.
5. Resuspended the pellet in 500 µl FBS and counted cells (4.5 million cells per tube).
6. Added 500 µl freezing medium and froze cells immediately in a Corning® CoolCell™ FTS30 at -80°C overnight.
7. Transferred cells to liquid nitrogen the next day.

#### Bait preparation

*For the following steps, a total cell number of 3 million cells per sample was assumed*.

SARS-CoV-2 receptor-binding domain (RBD) and S6P baits were freshly prepared the day before the staining experiment. The S6P bait contained trimeric SARS-CoV-2 Spike protein, which was biotinylated using the Avitag™ technology (ACROBiosystems, Cat. SPN-C82E9). The biotinylated protein was labeled at a 1:1 ratio with a fluorochrome-coupled streptavidin (BV480 or BV650). For staining, a 5 pmol bait solution for each dye in 25 µl per cell sample was prepared.

1. The fluorochrome-coupled streptavidin tube was centrifuged at 10,000 x g for 1 minutes at 4°C to spin-down any free dye particles.
2. 5 pmol BV480-coupled streptavidin were added to 5 pmol S6P protein directly in Brilliant staining buffer (total volume of 25 µl/cell sample).
3. 5 pmol BV650-coupled streptavidin to 5 pmol S6P protein were added directly to the Brilliant staining buffer (total volume of 25 µl/cell sample).
4. The baits were incubated overnight at 4°C, protected from light.

The RBD bait contains monomeric SARS-CoV-2 Spike RBD, biotinylated using the Avitag^TM^ technology (ACROBiosystems, Cat. SPD-C82E9). The biotinylated protein was labeled at a 4:1 ratio with a fluorochrome-coupled streptavidin (RB613 or RB744). For the staining, a 10 pmol bait solution for each dye in 25 µl/sample was prepared.

1. The fluorochrome-coupled streptavidin tube was centrifuged at 10,000 x g for 1 minutes at 4°C to spin-down any free dye particles.
2. In 4 increments with 20-minute[1] incubations between each addition, 2.5 pmol RB613-coupled streptavidin was added to 10 pmol RBD protein directly in Brilliant staining buffer (total volume of 25 µl/sample).
3. In 4 increments with 20-minute incubations between each addition, 2.5 pmol RB744-coupled streptavidin to 10 pmol RBD protein was added directly in Brilliant staining buffer (total volume of 25 µl/sample).
4. Baits were incubated overnight at 4°C, protected from light.

#### Staining of single cell suspensions

1. Removed cells from liquid nitrogen.
2. Thawed cells briefly in a 37°C water bath and transferred them immediately to a falcon containing 40 ml RPMI medium prewarmed to 37°C, allowing them to thaw completely.
3. Pelleted cells by centrifugation at 403 × g for 8 minutes at 4°C, aspirated the supernatant, and resuspended cells in 1 ml DPBS to remove any remaining proteins (e.g., FBS or bovine serum albumin) that could cause quenching of the free dye used for subsequent viability staining.
4. Counted cells and transferred 3 × 10⁶ cells per staining into an Eppendorf tube.
5. Washed cells twice by adding 500 µl of DPBS, then centrifuged tubes at 956 × g for 3 minutes at 4°C.
6. Resuspended cells in freshly prepared 100 µl Live-Dead dye solution.
7. Incubated for 15 minutes at RT in the dark.
8. Washed cells by adding 500 µl of Staining Buffer to remove unbound viability dye, centrifuged tubes at 956 × g for 3 minutes at 4°C, and discarded the supernatant.
9. Resuspended cells in 100 µl of Fc block (diluted 1:100 - purified anti-mouse CD16/CD32, clone 93) in Staining Buffer and added 5 µl CellBlox™ blocking buffer per 100 µl cell sample containing 10³ to 10⁸ cells (to block non-specific binding of NovaFluor labels, PE, and APC tandems observed with macrophages and monocytes).
10. Incubated for 15 minutes at 4°C, protected from light.
11. Washed cells by adding 500 µl of Staining Buffer to remove unbound viability dye, centrifuged tubes at 956 × g for 3 minutes at 4°C, and discarded the supernatant.
12. Resuspended cells in 100 µl of bait staining mix containing the correct final dilution of RBD and spike baits in Brilliant Staining Buffer.
13. Incubated for 1 hour at RT, protected from light.
14. Washed cells by adding 500 µl of Staining Buffer, centrifuged tubes at 956 × g for 3 minutes at 4°C, and discarded the supernatant.
15. Resuspended cells in 50 µl of antibody staining mix with the proper final dilutions of all antibodies diluted in Brilliant Staining Buffer.
16. Incubated for 15 minutes at 4°C, protected from light.
17. Washed cells twice by adding 500 µl of Staining Buffer per wash, centrifuged tubes at 956 × g for 3 minutes at 4°C, and discarded the supernatant.
18. Resuspended cells in 100 µl Staining Buffer and kept them in the dark at 4°C until analysis on the BD FACSymphony™ A5 SE.

### Staining workflow for mouse lymphoid panel

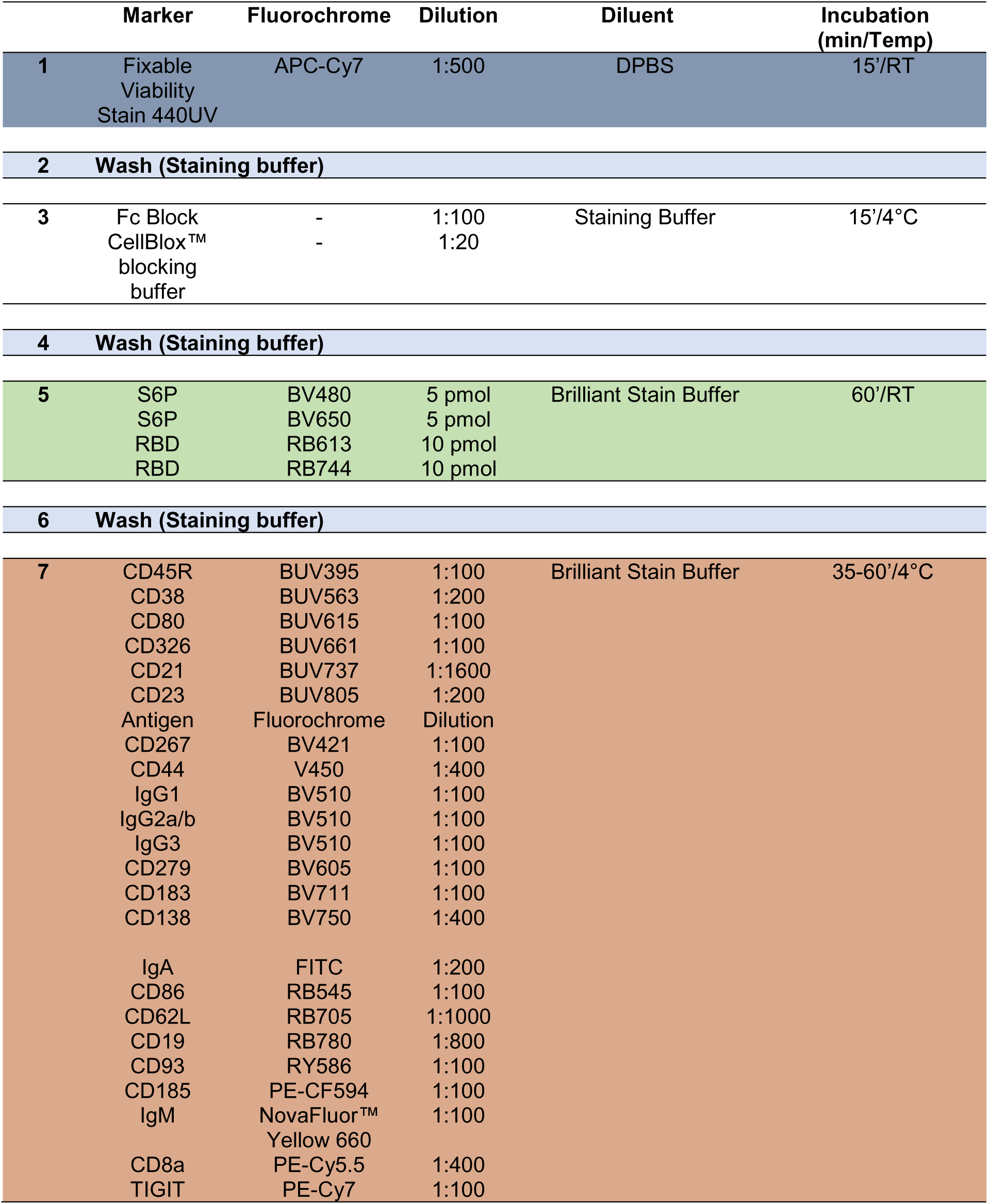
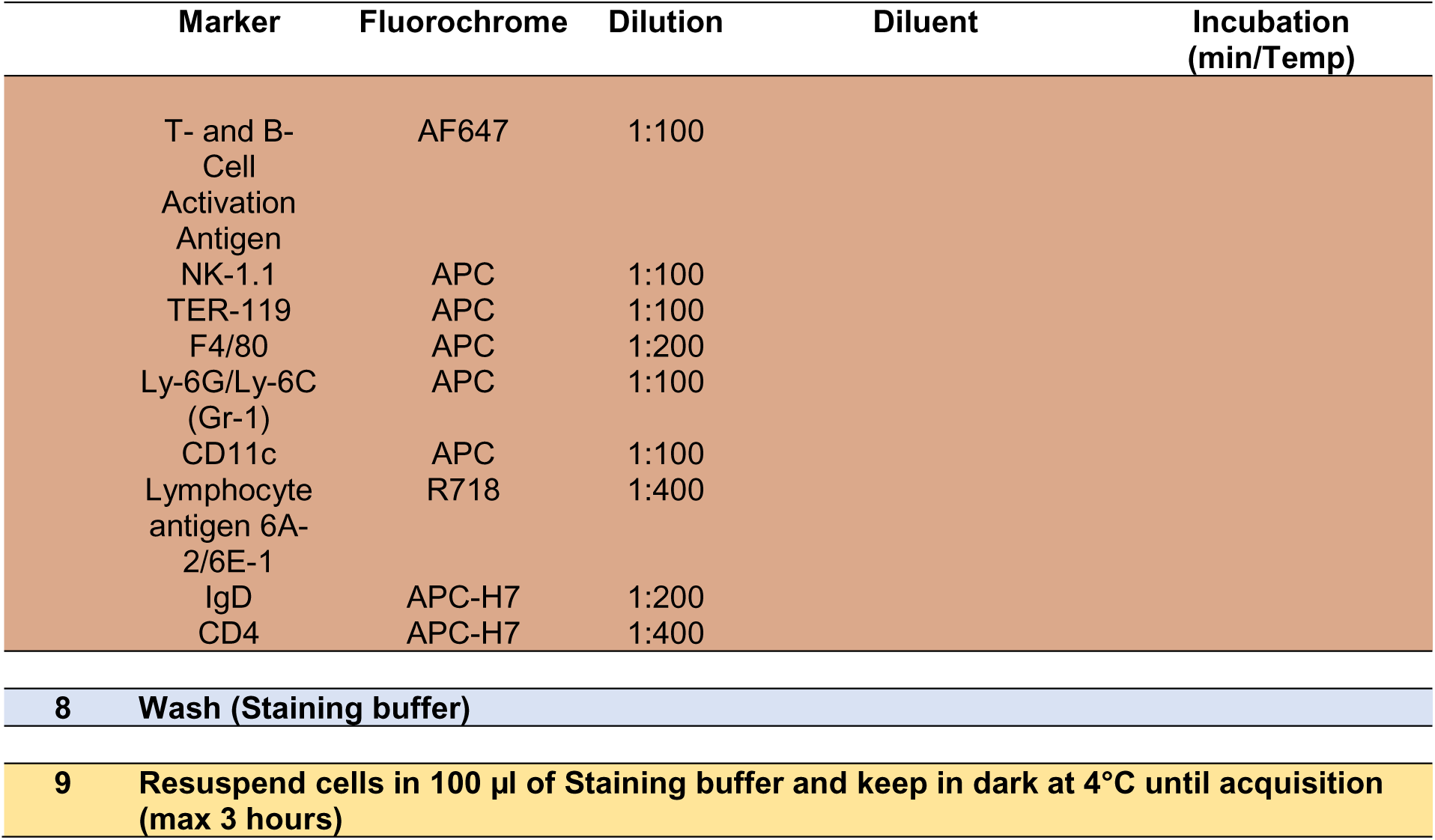

### Single Color Reference Control

Single stains of each antibody with either cells or beads were prepared. For unmixing, splenocytes, bone marrow-derived lymphocytes, and UltraComp eBeads™ Compensation Beads were used (Supplementary Table 4). We conducted an experiment in which we compared the quality of spectral unmixing of single-color reference controls prepared on either beads or cells. If beads yielded a comparable quality of spectral unmixing, they were chosen as single color reference control. For the baits, single-color reference controls based on antibodies against an abundantly expressed marker conjugated to the same fluorochrome from the same company (dummy approach) lead to satisfying results. Correct spectral unmixing was controlled with single stained cells. The separation of cells and populations was improved using autofluorescence unmixing.

#### Cell-based staining protocol for single-color reference controls

1. Up to 2×10^6^ cells per single stain were used. Cells were pelleted by centrifugation at 956 x g for 3 minutes at 4°C in Eppendorf tubes.
2. Resuspend cells in 100 µl Staining Buffer and add 5 µl CellBlox™ blocking buffer/ 100 µl cell single color reference control containing 10^3^ to 10^8^ cells (to block non-specific binding of NovaFluor labels, PE and APC tandems observed with macrophages and monocytes) where needed and incubate for 15 minutes at 4°C.
3. Wash cells by adding 500 µl of Staining buffer, centrifuge tubes at 956 x g for 3 minutes at 4°C and discard supernatant.
4. Resuspend cells in 100 µl of predetermined amount of antibody diluted in Brilliant staining buffer.
5. Incubate for 35-60 minutes at 4°C, protected from light.
6. Wash cells by adding 500 µl of Staining buffer, centrifuge tubes at 956 x g for 3 minutes at 4°C and discard supernatant.
7. Resuspend cells in 100 µl Staining buffer and keep in dark at 4°C until acquisition.
8. Collected 500 events within the negative and positive gates of interest.

#### Bead-based single color reference controls

The UltraComp eBeads™ Compensation Beads (Invitrogen, Cat. #01-2222-42) contained two bead populations. The positive bead population captured the fluorochrome-conjugated antibody used for cell staining, whereas the negative bead population did not bind the antibody.

1. One drop of UltraComp eBeads™ Compensation Beads was placed into an Eppendorf tube.
2. A predetermined amount of antibody diluted in Brilliant staining buffer (100 µl) was added.
3. Beads were incubated for 35–60 minutes at 4°C, protected from light.
4. Beads were washed by adding 500 µl staining buffer, centrifuged at 956 × g for 3 minutes at 4°C, and the supernatant was discarded.
5. Beads were resuspended in 100 µl staining buffer and stored in the dark at 4°C until acquisition.
6. 5000 events within the gate of interest were collected.

## Supplementary Information | Figures

**Supplementary Figure S1:**
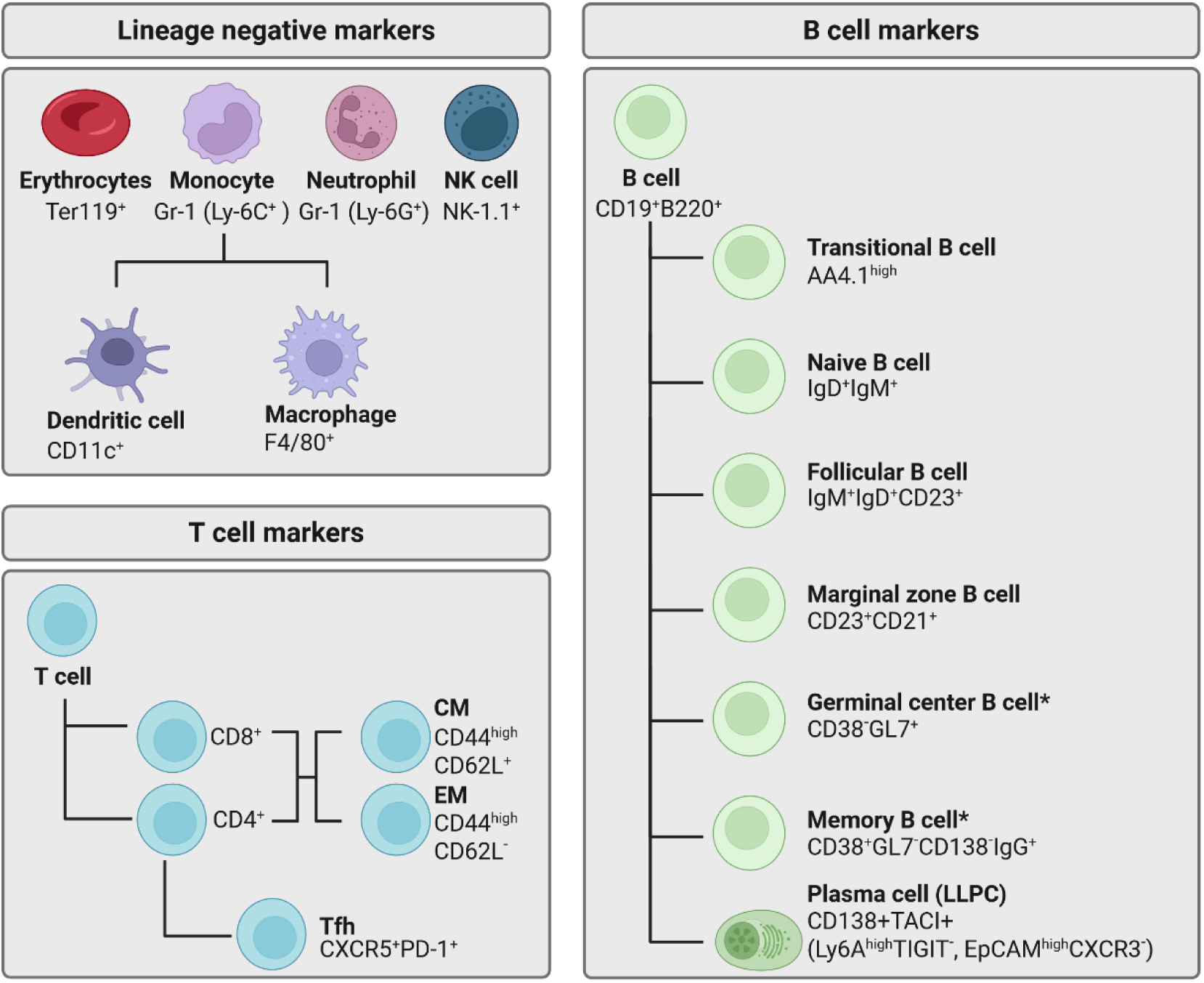
Overview of different lymphoid and myeloid subsets that can be discriminated with the panel. Phenotypic markers used to discriminate lymphoid cell populations including antigen-reactive B cells (subsets marked with an asterisk were analyzed) [1–7]. CM: Central memory, EM: Effector memory, Tfh: T follicular helper cell. Created in https://BioRender.com

**Supplementary Figure S2:**
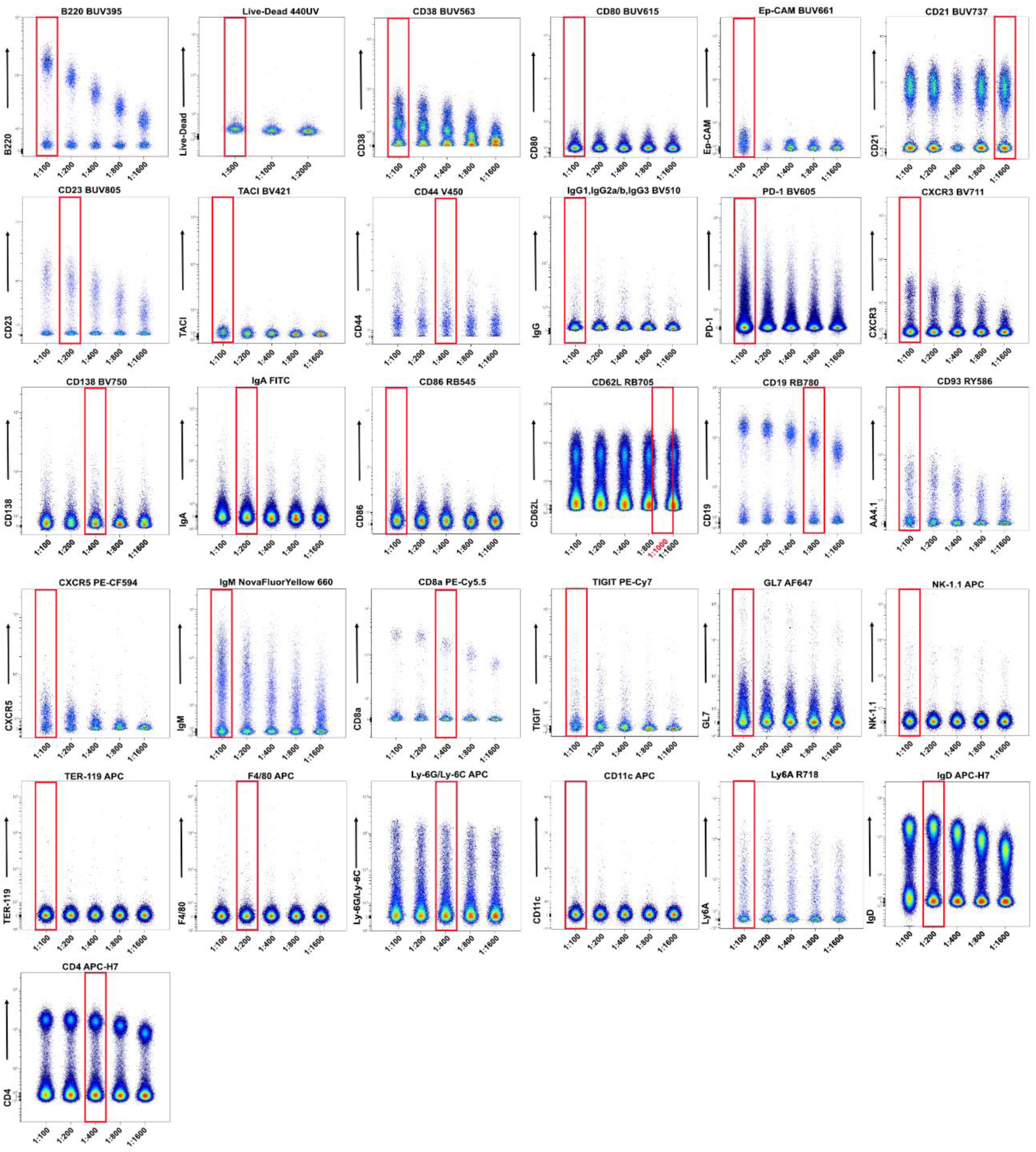
Antibody titrations for panel optimization. Antibody titration experiments were performed using splenocytes and bone marrow-derived lymphoid cells from 6–8-week-old C57BL/6 mice. Each antibody was tested in five 2-fold serial dilutions (indicated on the x-axis) in a final staining volume of 100 µL, with incubation for 35–60 minutes at 4°C in the dark. Individual FCS files were concatenated using OMIQ to visualize all dilutions in a single plot. Optimal antibody concentrations were defined as the lowest dilution providing maximal signal separation with minimal background staining. Red frames highlight dilutions used in the final panel.

**Supplementary Figure S3:**
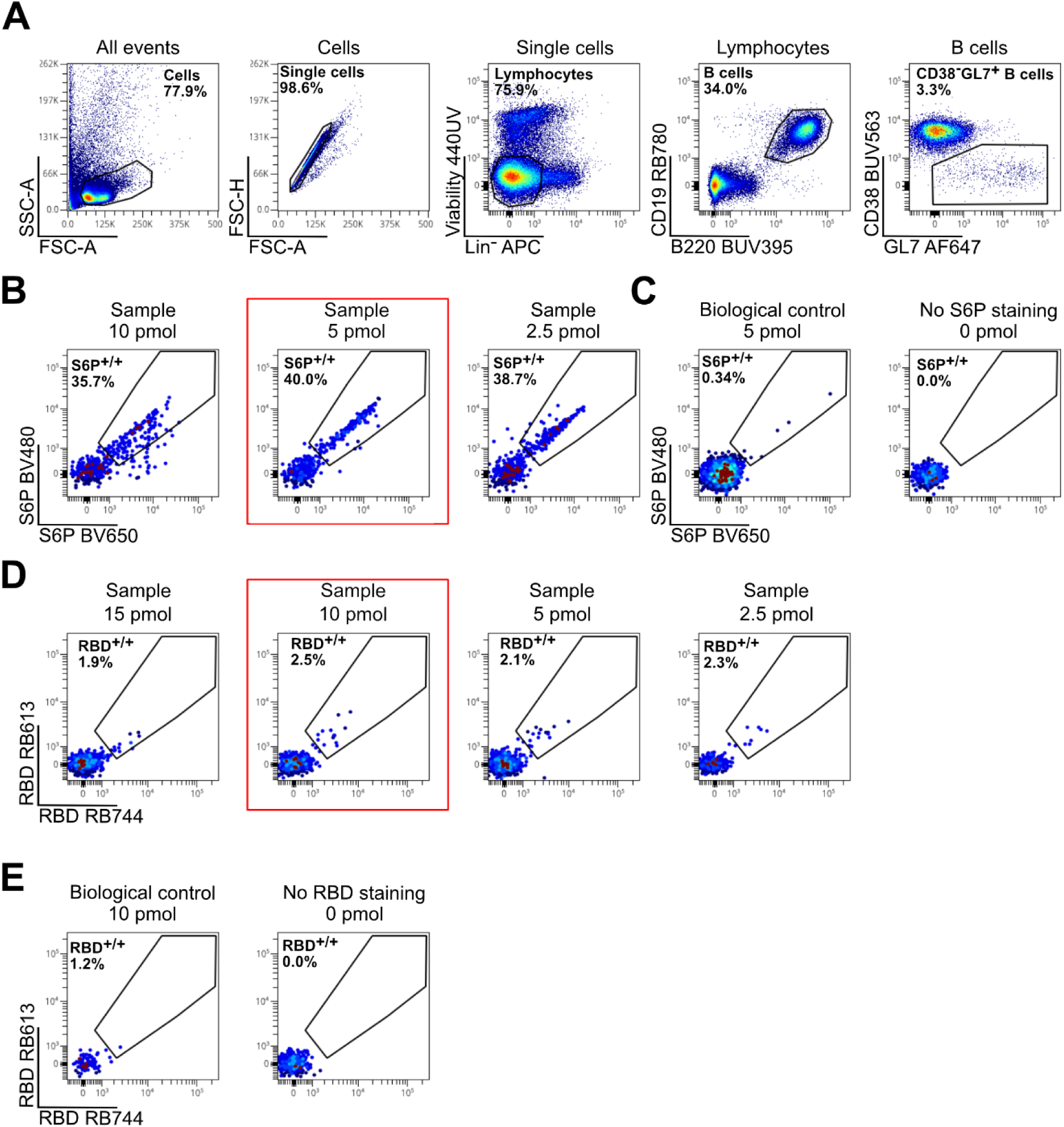
SARS-CoV-2 spike protein (S6P) and receptor binding domain (RBD) bait titration. Due to its large size (S6P, 419 kDa), the SARS-CoV-2 prefusion-stabilized spike protein is likely subject to steric hindrance that prevents tetramer formation when complexed with streptavidin (55 kDa), whose biotin-binding sites are separated by 20 Å (cis) and 35 Å (trans) [8]. Therefore, S6P and fluorochrome-conjugated streptavidin were combined at a 1:1 molar ratio to favor monomeric complexes. In contrast, the smaller receptor-binding domain (RBD, 28.2 kDa) likely forms tetramers and was thus used at a 4:1 molar ratio with streptavidin. Bait titration experiments were performed using lymph node-derived lymphoid cells from immunized mice to determine optimal concentrations for detecting antigen-reactive B cells. Cells were pre-incubated with serial bait concentrations (indicatedF above each plot) in 100 µL for 60 minutes at room temperature, followed by staining with the antibody mix in 50 µL for 35–60 minutes at 4°C, protected from light. Optimal bait concentrations were defined as the lowest amount yielding the best signal-to-noise separation with minimal background staining. Negative controls included cells from mice immunized with adjuvant only and spillover assessments of the antibody mix. Red frames highlight the bait concentrations selected for use in the final panel.

**Supplementary Figure S4:**
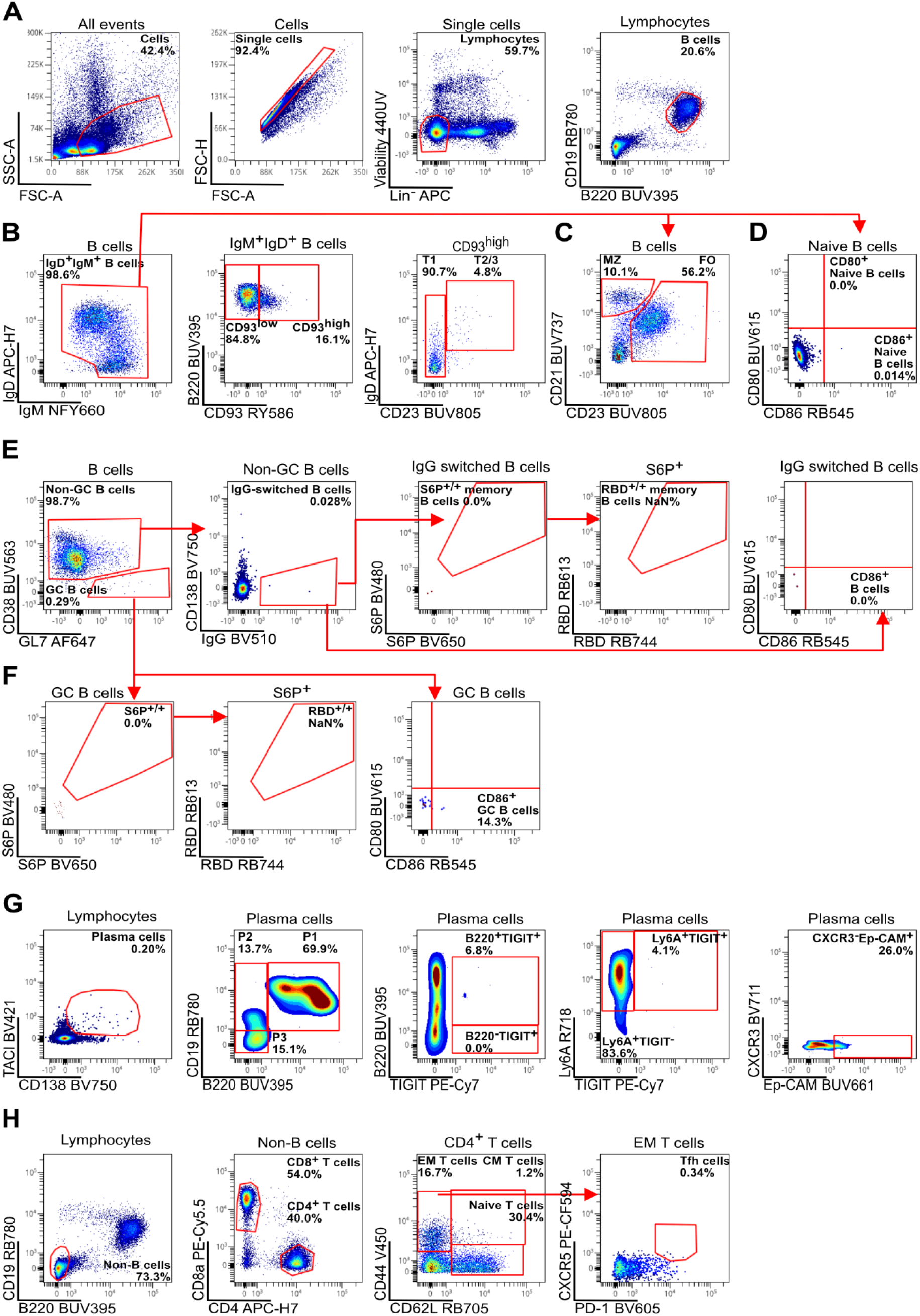
Gating strategy for murine spleen from a non-immunized mouse. A single cell suspension was generated from a spleen of a non-immunized mouse. The same gating as in Fig. 1 was applied. **(A)** Gating strategy for viable lymphocytes after exclusion of cellular debris, doublets, dead cells, and erythrocytes (TER^-^119^+^), monocytes and neutrophils (Ly-6G/Ly-6C^+^ (Gr-1^+^)) as well as macrophages (F4/80^+^), dendritic cells (CD11c^+^), and NK cells (NK-1.1^+^). B cells (CD19^+^B220^+^) were identified based on the surface expression of CD19 and B220. **(B)** From IgM^+^IgD^+^ B cells, CD93 (AA4.1)^high^ B cells were further gated to identify T1 (IgM^high^CD23^low^) and T2 (IgD^high^CD23^high^) stage transitional B cells. **(C)** Mature naive and marginal zone B cells were defined as CD23^high^CD21^+^ or CD23^low^CD21^high^.**(D)** IgM+IgD+ B cells were analyzed for expression of activation markers CD80 and CD86. **(E)** To identify memory B cells, plasma cells (PCs; CD138^+^) were excluded from non-germinal center (GC) B cells and further gated on class-switched B cells (IgG^+^). Among these IgG^+^class-switched B cells, antigen-reactive (S6P^+/+^, RBD^+/+^) cells were absent as well as activation marker (CD86^+^) expressing class-switched B cells. **(F)** GC B cells (CD38-GL7+) were identified based on the lack of CD38 expression. GC B cells were investigated for antigen-reactivity (S6P^+/+^, RBD^+/+^) as well as their activation status (CD86^+^). **(G)** PCs and plasmablasts (PBs) (Transmembrane activator and CAML interactor (TACI)^int^CD138+) were divided into subsets P1 – early dividing precursor PCs (B220^int^CD19^int^), P2 - early PCs (B220^low^CD19^int^), and P3 - mature resting PCs (B220^low^CD19^low^). Expression of TIGIT (and B220) was indicative of early and likely still proliferation P1 and P2 antibody secreting cells. CXCR3, Ep-CAM and Ly6A, markers were associated with IgG (Ep-CAM^high^CXCR3^-^) or IgA (Ly6A^high^TIGIT^-^) expression. **(H)** Among non-B cells, CD8^+^ and CD4^+^ T cells were discriminated. CD4^+^ T cells were separated into EM T cells (CD44^+^CD62L^-^), CM T cells (CD44^+^CD62L^+^), and naive T cells (CD44^-^CD62L^+^). EM T cells were further gated to identify Tfh cells (CXCR5^+^PD-1^+^).

**Supplementary Figure S5:**
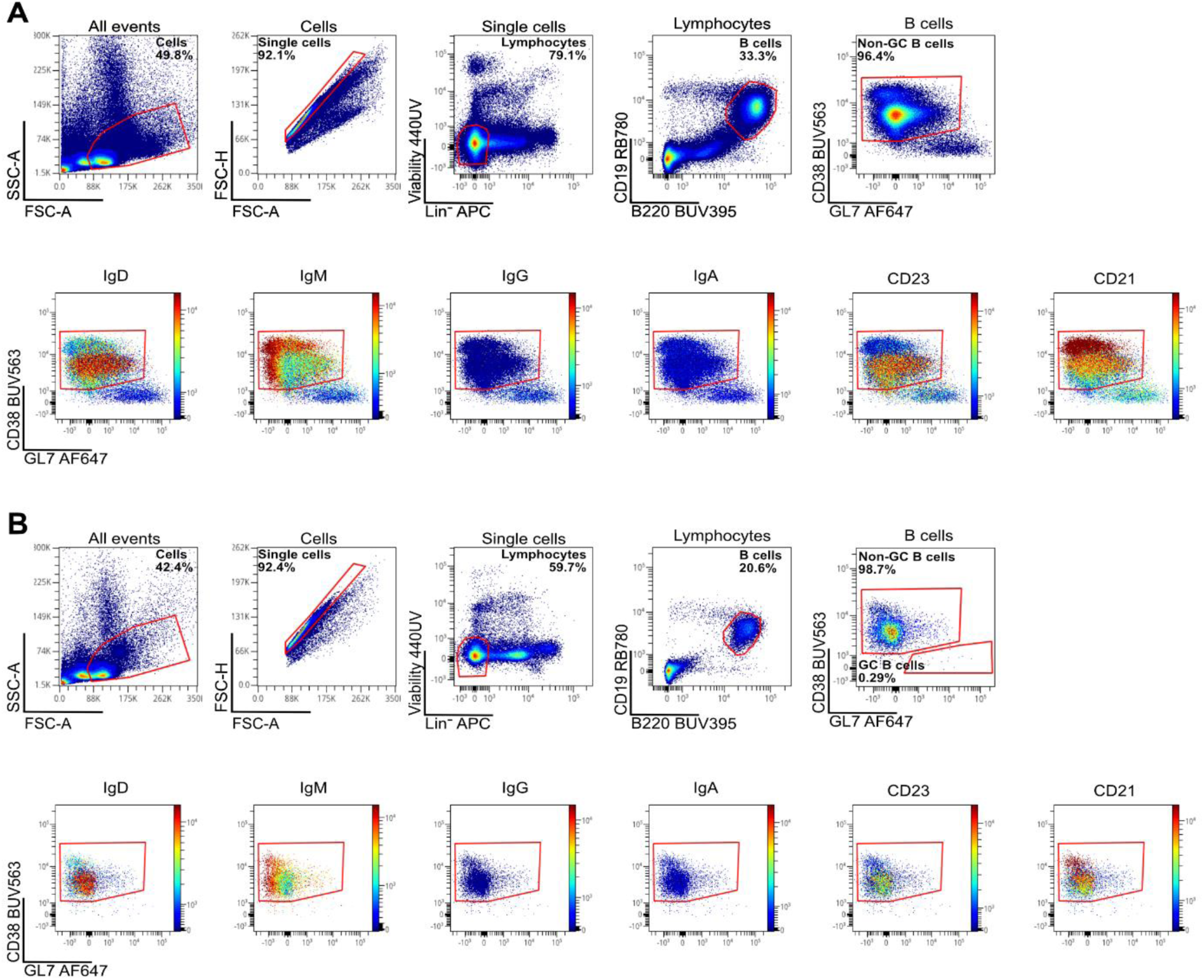
Gating strategy and phenotypic characterization of CD38^+^ non-GC B cell subsets in the spleen of immunized and non-immunized mice. (A) and (B) show representative data from an immunized **(A)** and a non-immunized **(B)** mouse. Gating strategy for viable lymphocytes after exclusion of cellular debris, doublets, dead cells, and erythrocytes (TER^-^119^+^), monocytes and neutrophils (Ly-6G/Ly-6C^+^ (Gr-1^+^)) as well as macrophages (F4/80^+^), dendritic cells (CD11c^+^), and NK cells (NK-1.1^+^). B cells (CD19^+^B220^+^) were identified using antibodies against CD19 and B220. B cells were further gated on non-GC (CD38^+^GL7^-^) B cells. Within the CD38^+^ population the expression of several markers was analyzed to identify marginal zone B cells (IgM^+^CD21^+^), mature naïve B cells (CD23^high^CD21^+^) and IgG+ class-switched B cells.

**Supplementary Figure S6:**
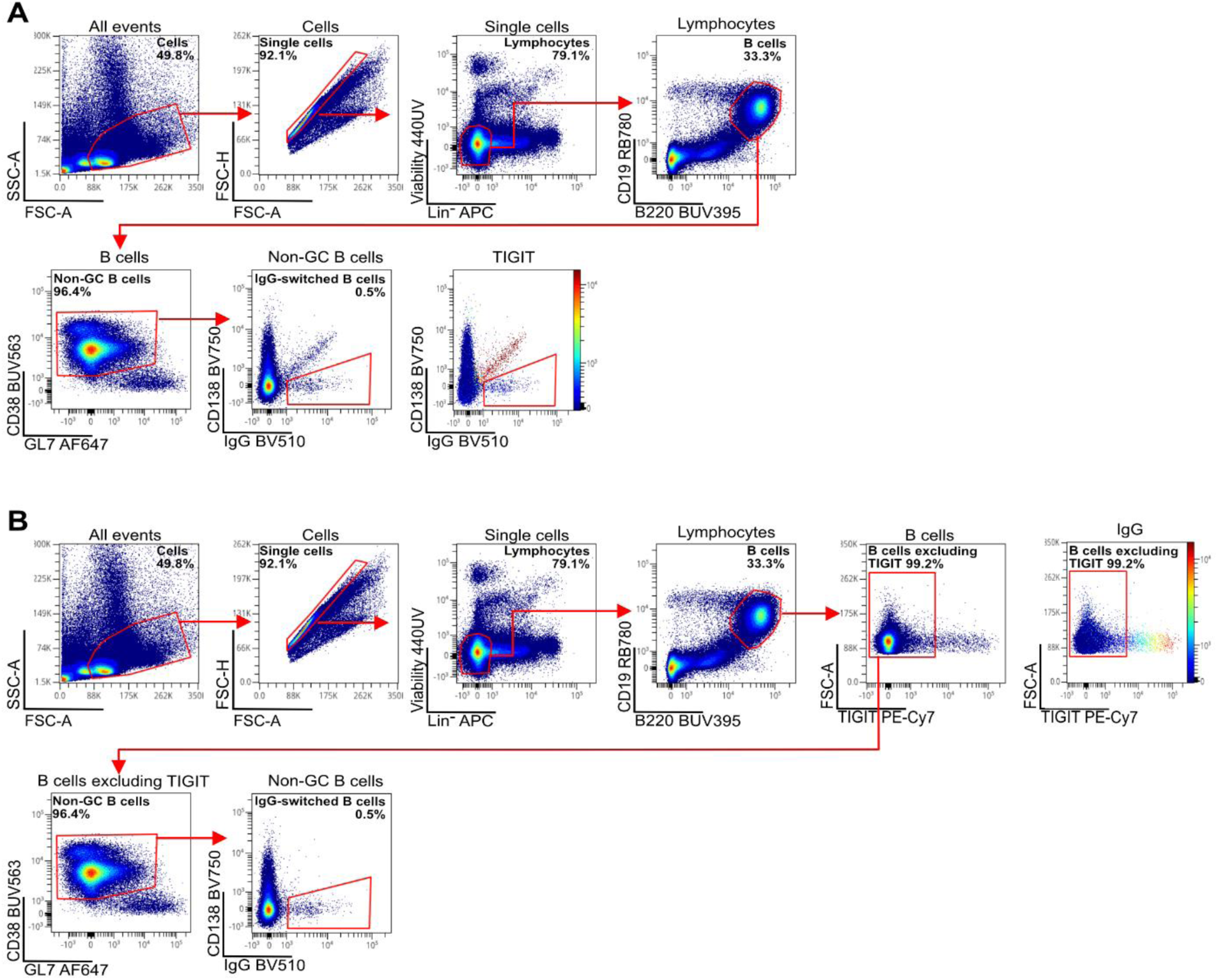
Adapted gating strategy for identifying IgG^+^ class-switched B cells. Gating strategy for viable lymphocytes after exclusion of cellular debris, doublets, dead cells, and erythrocytes (TER^-^119^+^), monocytes and neutrophils (Ly-6G/Ly-6C^+^ (Gr-1^+^)) as well as macrophages (F4/80^+^), dendritic cells (CD11c^+^), and NK cells (NK-1.1^+^). B cells (CD19^+^B220^+^) were identified using antibodies against CD19 and B220. **(A)** In the initial gating strategy, CD38^+^ non-GC B cells were further gated on IgG expression to identify class-switched B cells. However, this approach resulted in detectable spillover, likely caused by the anti-TIGIT antibody (a mouse IgG1 antibody) being bound by the anti-mouse IgG1 antibody. **(B)** To avoid this artifact, TIGIT^+^ B cells were excluded from the B cell population prior to gating on CD38^+^ non-GC B cells. This adapted strategy eliminated the false-positive IgG signal and allowed for clear identification of true IgG^+^ class-switched B cells.

**Supplementary Figure S7:**
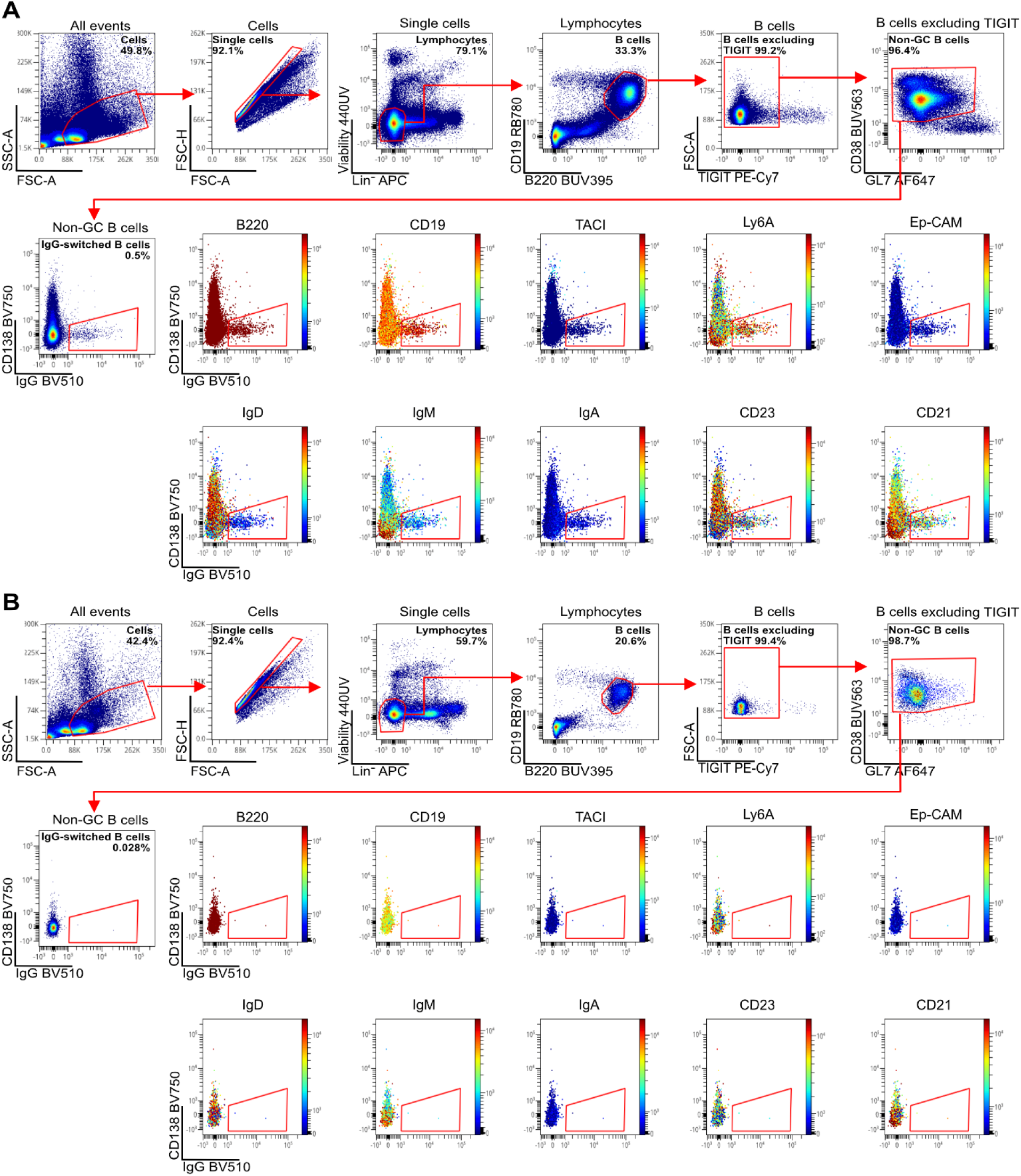
Gating on IgG^+^ class-switched B cells in the spleen. Gating strategy for viable lymphocytes after exclusion of cellular debris, doublets, dead cells, and erythrocytes (TER^-^119^+^), monocytes and neutrophils (Ly-6G/Ly-6C^+^ (Gr-1^+^)) as well as macrophages (F4/80^+^), dendritic cells (CD11c^+^), and NK cells (NK-1.1^+^). B cells (CD19^+^B220^+^) were identified using antibodies against CD19 and B220. To avoid false-positive staining due to spillover from the anti-TIGIT antibody (mouse IgG1 isotype), TIGIT^+^ B cells were excluded prior to further gating. **(A)** In a spleen sample from an immunized mouse, IgG^+^ class-switched B cells were clearly distinguishable from marginal zone B cells (IgM^+^CD21^+^), mature naïve B cells (CD23^high^CD21^+^), and PCs (CD138^+^TACI^+^) as shown by marker expression in the target population. **(B)** Equivalent analysis of a spleen sample from a non-immunized mouse, in which IgG^+^ class-switched B cells were largely absent.

**Supplementary Figure S8:**
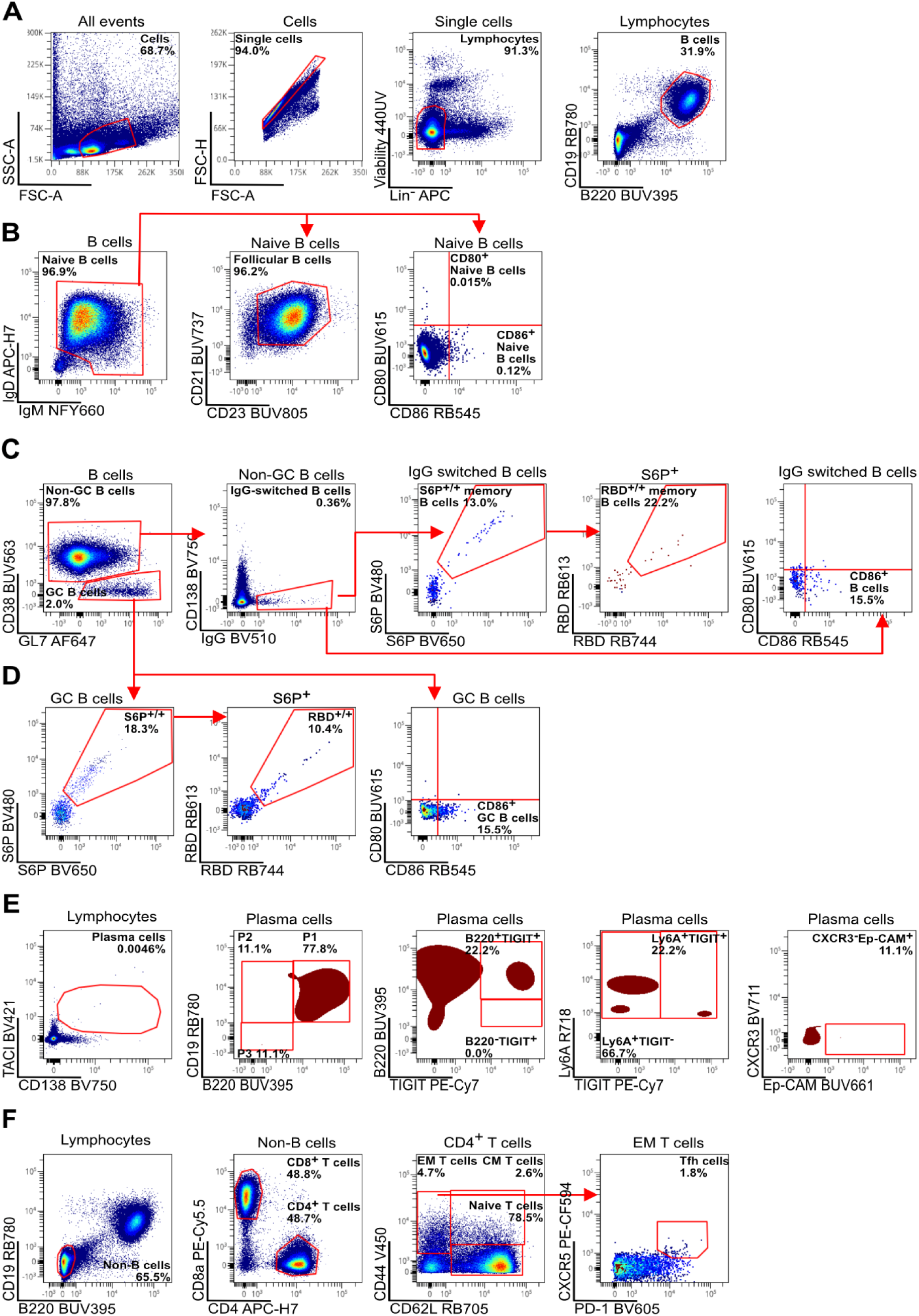
Gating strategy for murine lymph node samples. A single cell suspension was generated from pooled lymph nodes (inguinal, axial, and popliteal) of an immunized mouse 7 days after secondary immunization. **(A)** Gating strategy for viable lymphocytes after exclusion of cellular debris, doublets, dead cells, and erythrocytes (TER^-^119^+^), monocytes and neutrophils (Ly-6G/Ly-6C^+^ (Gr-1^+^)) as well as macrophages (F4/80^+^), dendritic cells (CD11c^+^), and NK cells (NK-1.1^+^). B cells (CD19^+^B220^+^) were identified based on the surface expression of CD19 and B220. **(B)** Lymph node samples mostly contained mature naïve B cells defined as IgM^low^IgD^high^CD23^high^CD21^+^. A small fraction of these cells showed elevated expression of the activation marker CD80 but not of CD86 compared to non-immunized mice. **(C)** To identify memory B cells, PCs (CD138^+^) were excluded from non-GC B cells and further gated on class-switched B cells (IgG^+^). Among IgG^+^ class-switched B cells, antigen-reactive (S6P^+/+^, RBD^+/+^) and activation marker (CD86^+^) expressing subsets were distinguished. **(D)** GC B cells (CD38-GL7+) were identified based on the absence of CD38 expression. GC B cells were investigated for antigen-reactivity (S6P^+/+^, RBD^+/+^) as well as their activation status (CD86^+^). **(E)** PCs and PBs (TACI^int^CD138+) were divided into subsets P1 – early dividing precursor PCs (B220^int^CD19^int^), P2 - early PCs (B220^low^CD19^int^), and P3 - mature resting PCs (B220^low^CD19^low^). Expression of TIGIT (and B220) was indicative of early and likely still proliferation P1 and P2 antibody secreting cells. CXCR3, Ep-CAM and Ly6A markers were associated with IgG (Ep-CAM^high^CXCR3^-^) or IgA (Ly6A^high^TIGIT^-^) expression. **(F)** Among non-B cells, CD8^+^ and CD4^+^ T cells were discriminated. CD4^+^ T cells were separated into EM (CD44^+^CD62L^-^), CM (CD44^+^CD62L^+^), and naive (CD44^-^CD62L^+^) T cells. EM T cells were further gated to identify Tfh cells (CXCR5^+^PD-1^+^).

**Supplementary Figure S9:**
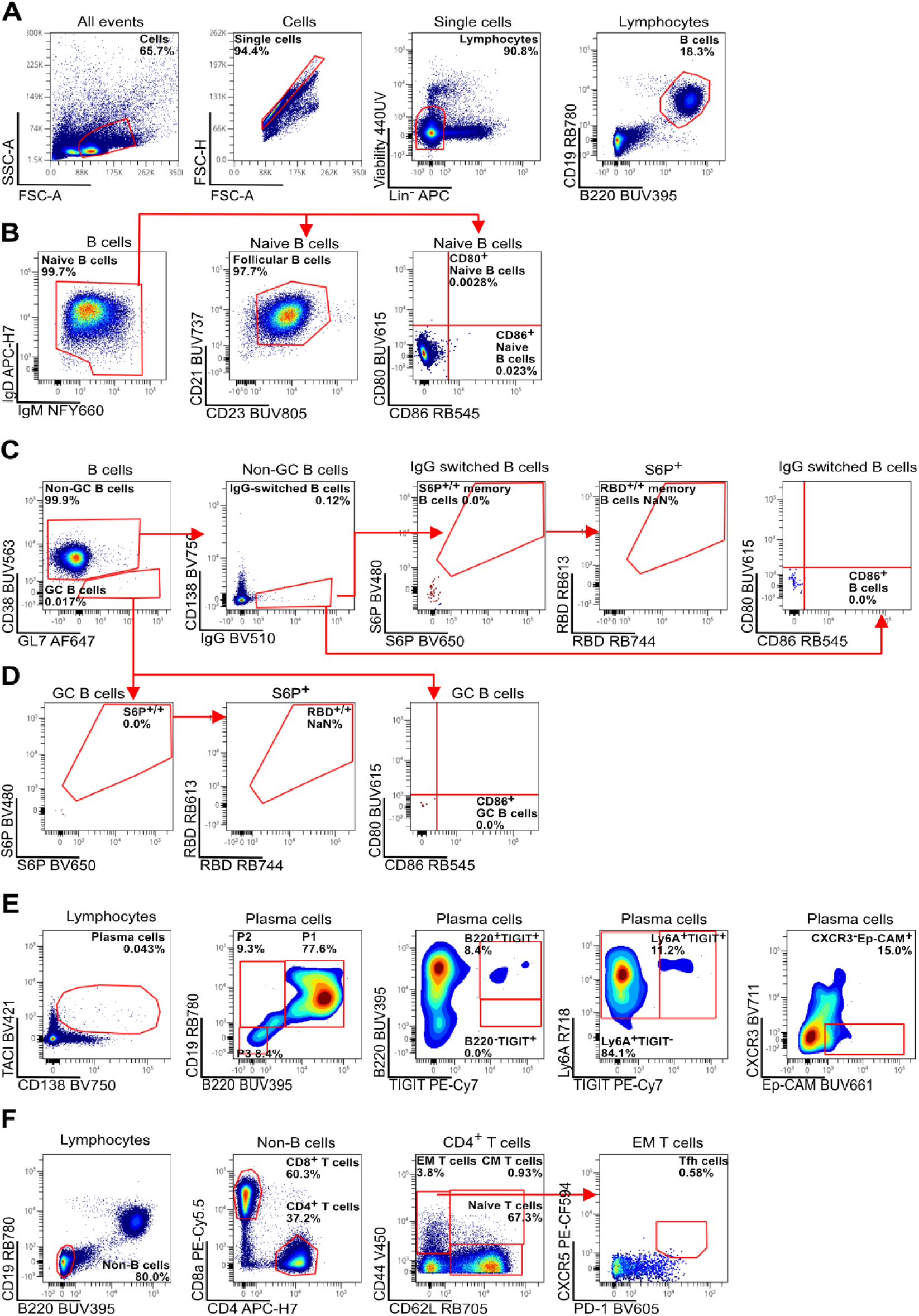
Gating strategy for murine lymph node samples. A single cell suspension was generated from pooled lymph node samples (inguinal, axial, and popliteal) of a non-immunized mouse. **(A)** Gating strategy for viable lymphocytes after exclusion of cellular debris, doublets, dead cells, and erythrocytes (TER^-^119^+^), monocytes and neutrophils (Ly-6G/Ly-6C^+^ (Gr-1^+^)) as well as macrophages (F4/80^+^), dendritic cells (CD11c^+^), and NK cells (NK-1.1^+^). B cells (CD19^+^B220^+^) were identified based on the surface expression of CD19 and B220. **(B)** Lymph node samples contained mostly mature naïve B cells defined as IgM^low^IgD^high^CD23^high^CD21^+^. These cells showed almost no expression of the activation markers CD80 and CD86 compared to immunized mice. **(C)** To identify memory B cells, PCs (CD138^+^) were excluded from non-GC B cells and further gated on class-switched B cells (IgG^+^). IgG^+^ class-switched B cells lacked antigen-reactivity (S6P^+/+^, RBD^+/+^) and activation marker (CD86^+^) expression. **(D)** GC B cells (CD38-GL7+) were identified based on the absence of CD38 expression. GC B cells were investigated for antigen-reactivity (S6P^+/+^, RBD^+/+^) and activation status (CD86^+^). **(E)** PCs and PBs (TACI^int^CD138+) were divided into subsets P1 – early dividing precursor PCs (B220^int^CD19^int^), P2 - early PCs (B220^low^CD19^int^), and P3 - mature resting PCs (B220^low^CD19^low^). Expression of TIGIT (and B220) was indicative of early and likely still proliferation P1 and P2 antibody secreting cells. CXCR3, Ep-CAM and Ly6A, markers were associated with IgG (Ep-CAM^high^CXCR3^-^) or IgA (Ly6A^high^TIGIT^-^) expression. **(F)** Among non-B cells, CD8^+^ and CD4^+^T cells were discriminated. CD4^+^ T cells were separated into EM (CD44^+^CD62L^-^), CM (CD44^+^CD62L^+^), and naive (CD44^-^CD62L^+^) T cells. EM T cells were further gated to identify Tfh cells (CXCR5^+^PD-1^+^).

**Supplementary Figure S10:**
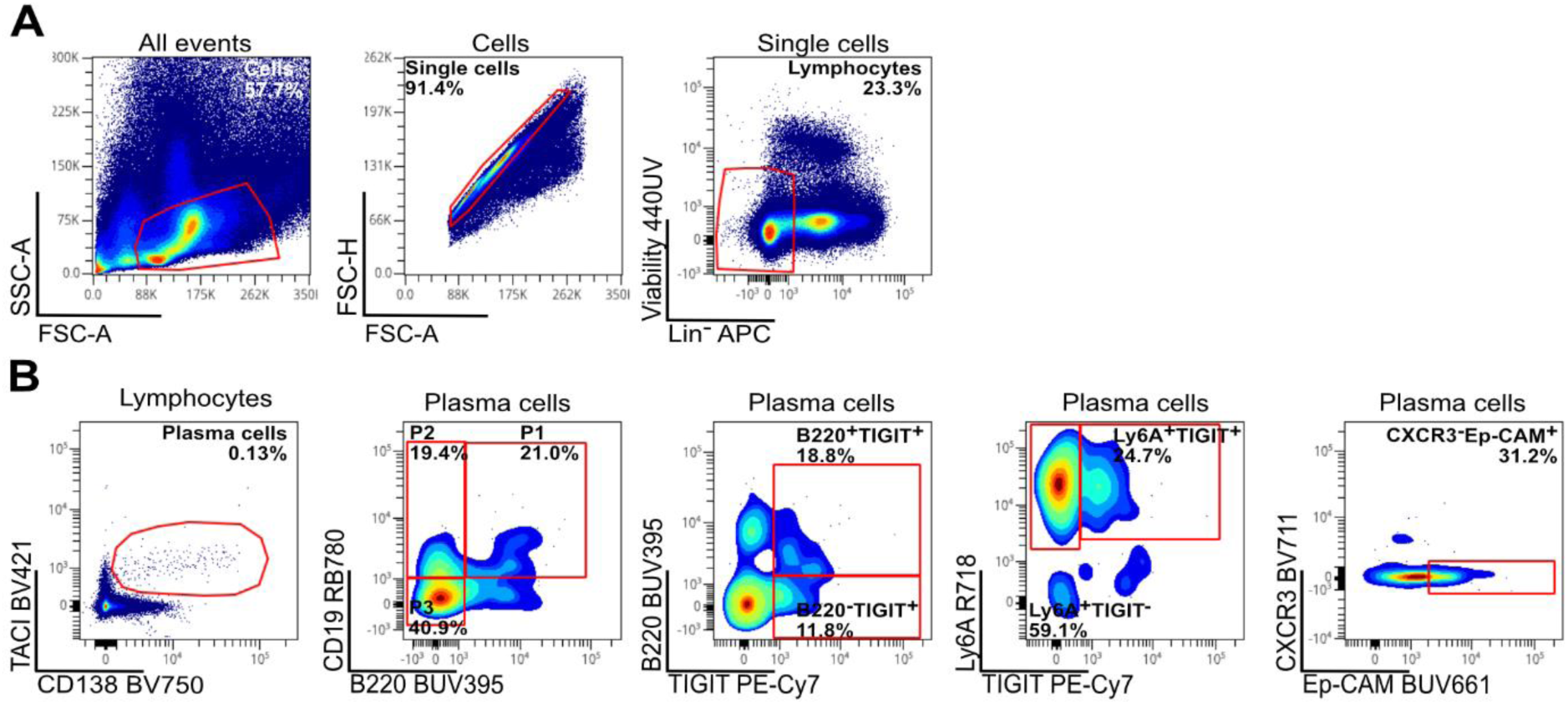
Gating strategy for murine bone-marrow PCs. Single cell suspension from bone marrow of a non-immunized mouse. **(A)** Gating strategy for viable lymphocytes after exclusion of cellular debris, doublets, dead cells, and erythrocytes (TER^-^119^+^), monocytes and neutrophils (Ly-6G/Ly-6C^+^ (Gr-1^+^)) as well as macrophages (F4/80^+^), dendritic cells (CD11c^+^), and NK cells (NK-1.1^+^). **(B)** PCs and PBs (TACI^int^CD138+) were divided into subsets P1 – early dividing precursor PCs (B220^int^CD19^int^), P2 - early PCs (B220^low^CD19^int^), and P3 - mature resting PCs (B220^low^CD19^low^). Expression of TIGIT (and B220) was indicative of early and likely still proliferation P1 and P2 antibody secreting cells. CXCR3, Ep-CAM and Ly6A, markers were associated with IgG (Ep-CAM^high^CXCR3^-^) or IgA (Ly6A^high^TIGIT^-^) expression.

**Supplementary Figure S11:**
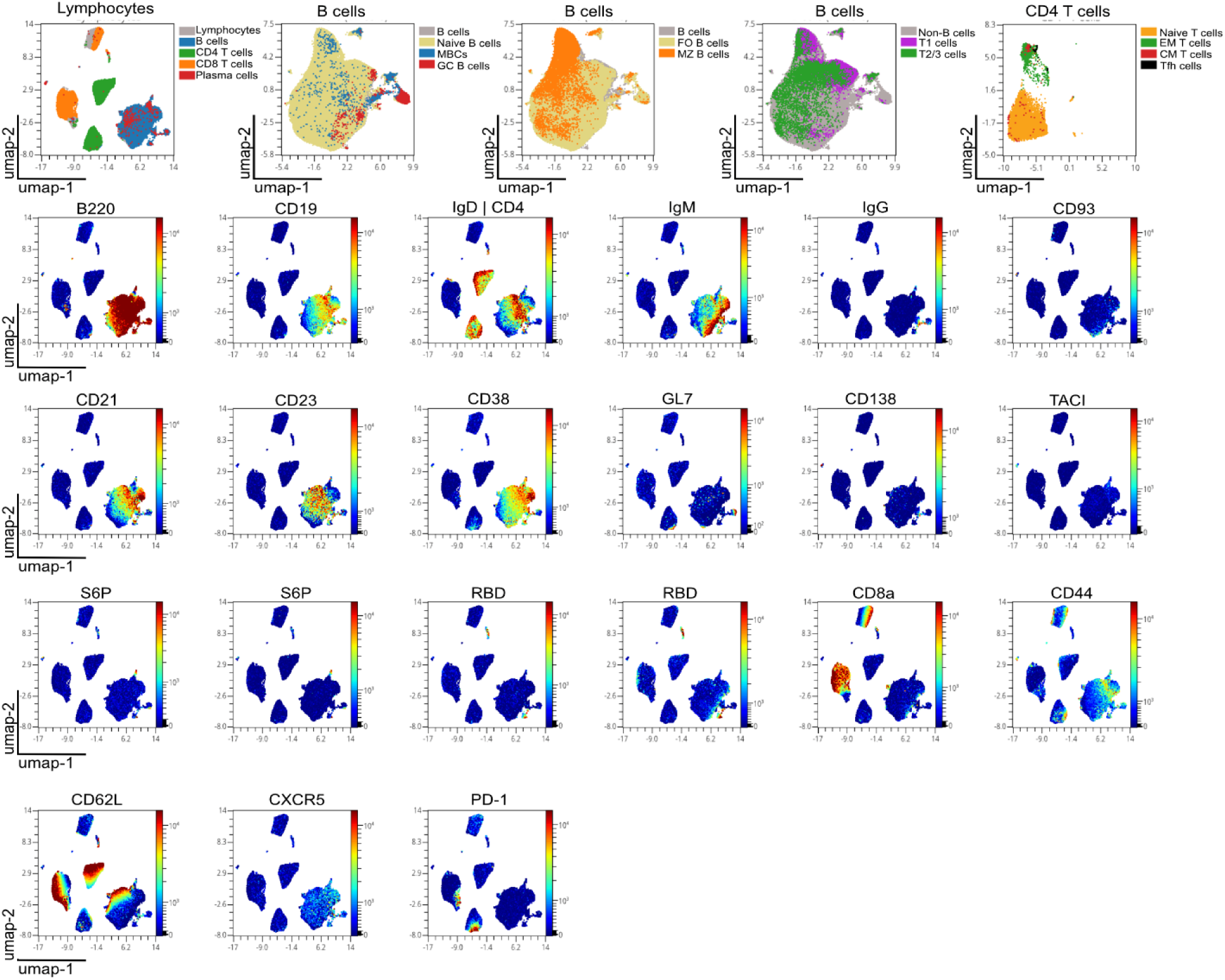
Uniform Manifold Approximation and Projection (UMAP) analysis of data collected from a spleen sample of an immunized mouse 7 days after secondary immunization. For the dimensionality reduction approach, cellular debris, doublets, dead cells, and erythrocytes (TER-119+), monocytes and neutrophils (Ly-6G/Ly-6C+ (Gr-1+)) as well as macrophages (F4/80+), dendritic cells (CD11c+), and NK cells (NK-1.1+) were excluded. UMAP was applied on the lymphocytes gate using OMIQ (Nearest Neighbors = 50, Minimum Distance = 0.1, Metric = Euclidean, Components = 2). Overlay of the four main populations B cells, CD4 T cells, CD8 T cells, and PCs. Overlay of distinct B cell and T cell populations shown in several plots for better resolution.

**Supplementary Figure S12:**
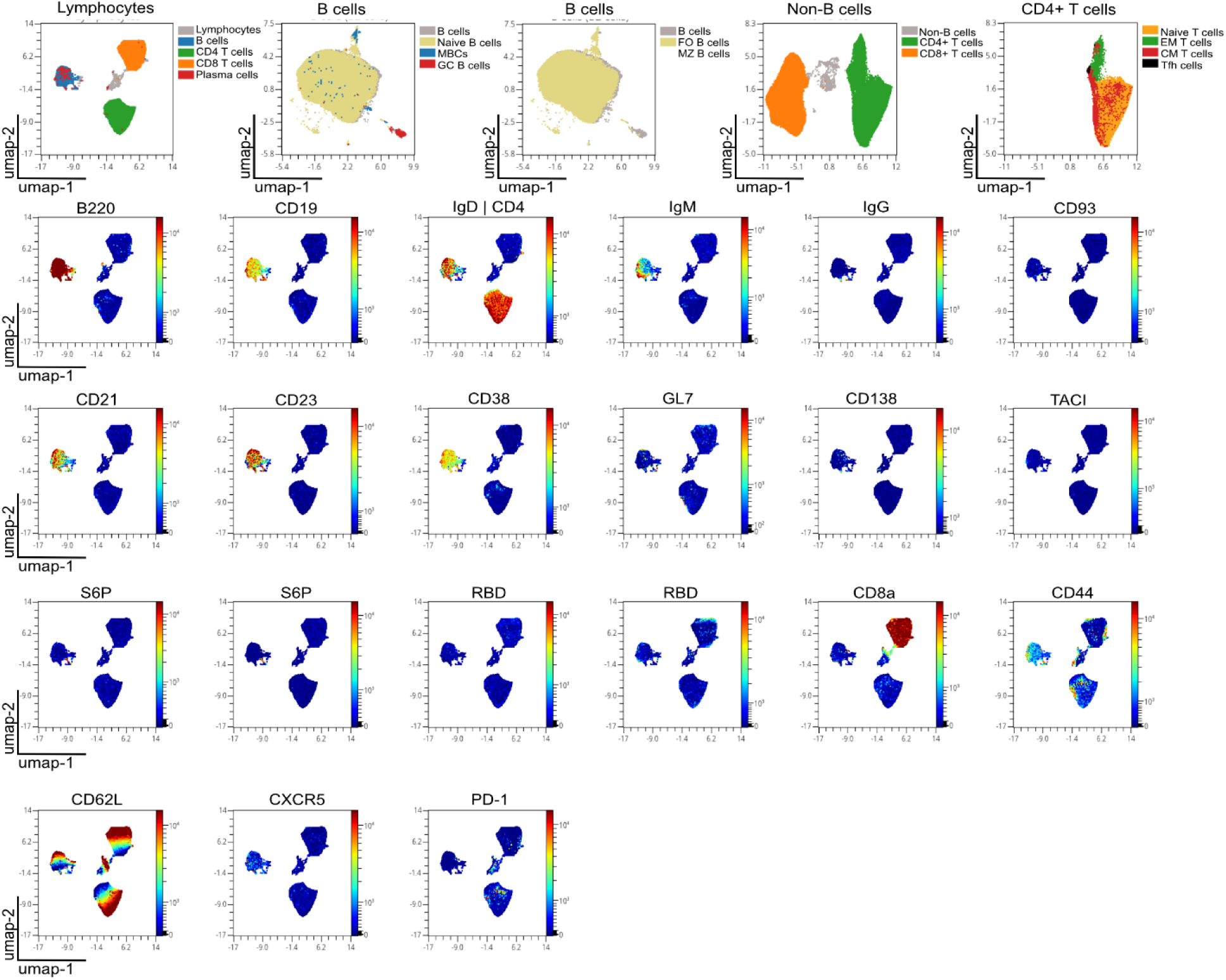
UMAP analysis of data collected from pooled lymph node samples (inguinal, axial, and popliteal) of an immunized mouse 7 days after secondary immunization. For the dimensionality reduction approach, cellular debris, doublets, dead cells, and erythrocytes (TER-119+), monocytes and neutrophils (Ly-6G/Ly-6C+ (Gr-1+)) as well as macrophages (F4/80+), dendritic cells (CD11c+), and NK cells (NK-1.1+) were excluded. UMAP was applied on the lymphocytes gate using OMIQ (Nearest Neighbors = 50, Minimum Distance = 0.1, Metric = Euclidean, Components = 2). Overlay of the four main populations B cells, CD4 T cells, CD8 T cells, and PCs. Overlay of distinct B cell and T cell populations shown in several plots for better resolution.

## Supplementary Information | Tables

**Supplementary Table 1:** Antibody clones and fluorochromes.

| Target specificity | Alternative Name | Clone | Fluorochrome | Purpose |
| --- | --- | --- | --- | --- |
| CD45R | B220 | RA3-6B2 | BUV395 | B cells |
| Live/Dead |  | N/A | Fixable Viability Stain 440UV | Viable cells |
| CD38 |  | 90 | BUV563 | B cells |
| CD80 |  | 16-10A1 | BUV615 | Activation |
| CD326 | Ep-CAM | G8.8 | BUV661 | B cells |
| CD21 |  | 7G6 | BUV737 | B cells |
| CD23 |  | B3B4 | BUV805 | B cells |
| CD267 | TACI | 8F10 | BV421 | B cells |
| CD44 |  | IM7 | V450 | T cells |
| S6P |  | N/A | BV480 | Antigen-reactivity |
| IgG1 |  | RMG1-1 | BV510 | B cells |
| IgG2a/b |  | R2-40 | BV510 | B cells |
| IgG3 |  | R40-82 | BV510 | B cells |
| CD279 | PD-1 | 29F.1A12 | BV605 | T cells |
| S6P |  | N/A | BV650 | Antigen-reactivity |
| CD183 | CXCR3 | CXCR3-173 | BV711 | B cells |
| CD138 | Synd-1 | 281-2 | BV750 | B cells |
| IgA |  | C10-3 | FITC | B cells |
| CD86 |  | PO3 | RB545 | Activation |
| RBD |  | N/A | RB613 | Antigen-reactivity |
| CD62L |  | MEL-14 | RB705 | T cells |
| RBD |  | N/A | RB744 | Antigen-reactivity |
| CD19 |  | 1D3 | RB780 | B cells |
| CD93 | AA4.1 | AA4.1 | RY586 | B cells |
| CD185 | CXCR5 | 2G8 | PE-CF594 | T cells |
| IgM |  | II/41 | NovaFluor™ Yellow 660 | B cells |
| CD8a |  | 53-6.7 | PE-Cy5.5 | T cells |
| TIGIT |  | 1G9 | PE-Cy7 | B cells |
| T- and B- Cell Activation Antigen | GL7 | GL7 | AF647 | B cells |
| NK-1.1 |  | PK136 | APC | Lineage negative |
| TER-119 |  | TER-119 | APC | Lineage negative |
| F4/80 |  | T45-2342 | APC | Lineage negative |
| Ly-6G/Ly-6C (Gr-1) |  | RB6-8C5 | APC | Lineage negative |
| CD11c |  | HL3 | APC | Lineage negative |
| Lymphocyte antigen 6A-2/6E-1 | Ly6A | D7 | R718 | B cells |
| IgD |  | 11-26c.2a | APC-H7 | B cells |
| CD4 |  | GK1.5 | APC-H7 | T cells |

**Supplementary Table 2:** Configuration of the full visible spectrum cytometer BD FACSymphony™ A5 SE with five lasers and 49 detectors.

| Wavelength (nm)/laser | Laser power (mW) | Detector* | Longpass optical filter | Bandpass (nm) optical filter | Fluorochrome (Peak intensity) |
| --- | --- | --- | --- | --- | --- |

| UV laser |  |  |  |  |  |
| --- | --- | --- | --- | --- | --- |
| 355/UV | 65 | A | 765LP | 809/82 | BUV805 |
|  |  | B | 704LP | 736/64 | BUV737 |
|  |  | C | 675LP | 695/40 |  |
|  |  | D | 645LP | 660/30 | BUV661 |
|  |  | E | 595LP | 610/30 | BUV615 |
|  |  | F | 570LP | 585/30 | BUV563 |
|  |  | G | 535LP | 540/20 | Autofluorescence measurement |
|  |  | H | 495LP | 515/30 |  |
|  |  | I | 425LP | 446/67 | Fixable Viability Stain 440UV |
|  |  | J | 365LP | 379/34 | BUV395 |

| Violet laser |  |  |  |  |  |
| --- | --- | --- | --- | --- | --- |
| 405/Violet | 200 | A | 810LP | 845/70 |  |
|  |  | B | 765LP | 785/50 |  |
|  |  | C | 730LP | 750/40 | BV750 |
|  |  | D | 690LP | 710/40 | BV711 |
|  |  | E | 665LP | 680/30 |  |
|  |  | F | 645LP | 660/30 | BV650 |
|  |  | G | 605LP | 615/25 |  |
|  |  | H | 585LP | 595/30 | BV605 |
|  |  | I | 570LP | 576/20 |  |
|  |  | J | 530LP | 540/20 |  |
|  |  | K | 495LP | 510/40 | BV510 |
|  |  | L | 465LP | 470/15 | BV480 |
|  |  | M | 430LP | 450/40 | V450 |
|  |  | N | 415LP | 431/28 | BV421 |

| Blue laser |  |  |  |  |  |
| --- | --- | --- | --- | --- | --- |
| 488/Blue | 150 | A | 770LP | 810/79 | RB780 |
|  |  | B | 724LP | 750/60 | RB744 |
|  |  | C | 685LP | 710/50 | RB705 |
|  |  | D | 665LP | 675/20 |  |
|  |  | E | 645LP | 660/30 |  |
|  |  | F | 585LP | 602/40 | RB613 |
|  |  | G | 570LP | 576/20 | RB545 |
|  |  | H | 520LP | 537/32 |  |
|  |  | I | 500LP | 510/20 | FITC |
|  |  | J | - | 488/10 | SSC |

| Wavelength<br>(nm)/Laser | Laser power<br>(mW) | Detector | Longpass | Bandpass<br>(nm) | Fluorochrome<br>(Peak<br>intensity) |
| --- | --- | --- | --- | --- | --- |

| Yellow Green laser |  |  |  |  |  |
| --- | --- | --- | --- | --- | --- |
| 561/Yellow-green | 150 | A | 800LP | 825/49 |  |
|  |  | B | 750LP | 780/60 | PE-Cy7 |
|  |  | C | 735LP | 750/40 |  |
|  |  | D | 699LP | 730/50 |  |
|  |  | E | 680LP | 695/40 | PeCy5.5 |
|  |  | F | 665LP | 670/20 |  |
|  |  | G | 645LP | 660/30 | NovaFluor™<br>Yellow 660 |
|  |  | H | 595LP | 602/40 | PE-CF594 |
|  |  | I | 570LP | 585/30 | RY586 |

| Red laser |  |  |  |  |  |
| --- | --- | --- | --- | --- | --- |
| 640/Red | 140 | A | 750LP | 780/60 | APC-H7 |
|  |  | B | 720LP | 730/50 | R718 |
|  |  | C | 699LP | 710/25 |  |
|  |  | D | 680LP | 680/30 | APC |
|  |  | E | 665LP | 675/20 | AF647 |
|  |  | F | 645LP | 662/11 |  |

**Supplementary Table 3:**
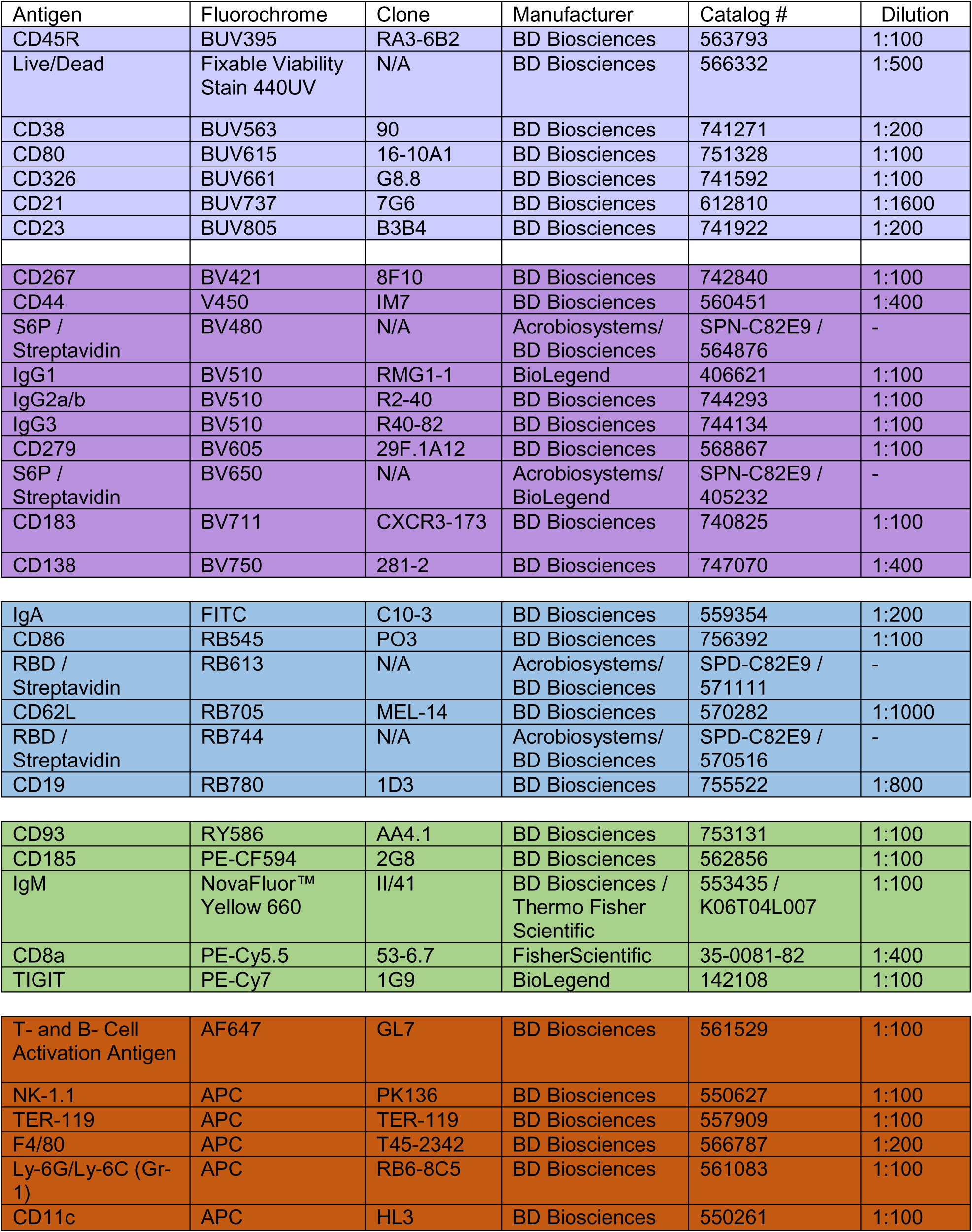
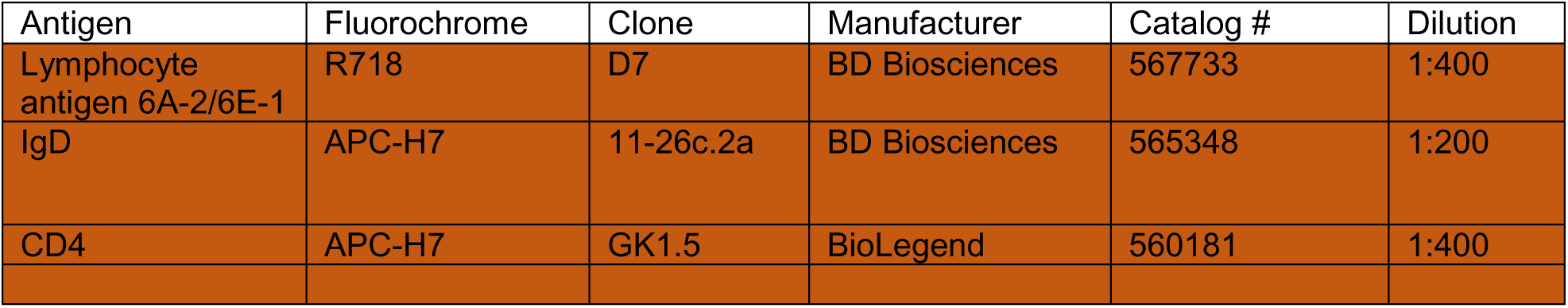
Antibody list.

| Antigen | Fluorochrome | Clone | Manufacturer | Catalog # | Dilution |
| --- | --- | --- | --- | --- | --- |
| CD45R | BUV395 | RA3-6B2 | BD Biosciences | 563793 | 1:100 |
| Live/Dead | Fixable Viability Stain 440UV | N/A | BD Biosciences | 566332 | 1:500 |
| CD38 | BUV563 | 90 | BD Biosciences | 741271 | 1:200 |
| CD80 | BUV615 | 16-10A1 | BD Biosciences | 751328 | 1:100 |
| CD326 | BUV661 | G8.8 | BD Biosciences | 741592 | 1:100 |
| CD21 | BUV737 | 7G6 | BD Biosciences | 612810 | 1:1600 |
| CD23 | BUV805 | B3B4 | BD Biosciences | 741922 | 1:200 |
| CD267 | BV421 | 8F10 | BD Biosciences | 742840 | 1:100 |
| CD44 | V450 | IM7 | BD Biosciences | 560451 | 1:400 |
| S6P / Streptavidin | BV480 | N/A | Acrobiosystems/ BD Biosciences | SPN-C82E9 / 564876 | - |
| IgG1 | BV510 | RMG1-1 | BioLegend | 406621 | 1:100 |
| IgG2a/b | BV510 | R2-40 | BD Biosciences | 744293 | 1:100 |
| IgG3 | BV510 | R40-82 | BD Biosciences | 744134 | 1:100 |
| CD279 | BV605 | 29F.1A12 | BD Biosciences | 568867 | 1:100 |
| S6P / Streptavidin | BV650 | N/A | Acrobiosystems/ BioLegend | SPN-C82E9 / 405232 | - |
| CD183 | BV711 | CXCR3-173 | BD Biosciences | 740825 | 1:100 |
| CD138 | BV750 | 281-2 | BD Biosciences | 747070 | 1:400 |
| IgA | FITC | C10-3 | BD Biosciences | 559354 | 1:200 |
| CD86 | RB545 | PO3 | BD Biosciences | 756392 | 1:100 |
| RBD / Streptavidin | RB613 | N/A | Acrobiosystems/ BD Biosciences | SPD-C82E9 / 571111 | - |
| CD62L | RB705 | MEL-14 | BD Biosciences | 570282 | 1:1000 |
| RBD / Streptavidin | RB744 | N/A | Acrobiosystems/ BD Biosciences | SPD-C82E9 / 570516 | - |
| CD19 | RB780 | 1D3 | BD Biosciences | 755522 | 1:800 |
| CD93 | RY586 | AA4.1 | BD Biosciences | 753131 | 1:100 |
| CD185 | PE-CF594 | 2G8 | BD Biosciences | 562856 | 1:100 |
| IgM | NovaFluor™ Yellow 660 | II/41 | BD Biosciences / Thermo Fisher Scientific | 553435 / K06T04L007 | 1:100 |
| CD8a | PE-Cy5.5 | 53-6.7 | FisherScientific | 35-0081-82 | 1:400 |
| TIGIT | PE-Cy7 | 1G9 | BioLegend | 142108 | 1:100 |
| T- and B- Cell Activation Antigen | AF647 | GL7 | BD Biosciences | 561529 | 1:100 |
| NK-1.1 | APC | PK136 | BD Biosciences | 550627 | 1:100 |
| TER-119 | APC | TER-119 | BD Biosciences | 557909 | 1:100 |
| F4/80 | APC | T45-2342 | BD Biosciences | 566787 | 1:200 |
| Ly-6G/Ly-6C (Gr-1) | APC | RB6-8C5 | BD Biosciences | 561083 | 1:100 |
| CD11c | APC | HL3 | BD Biosciences | 550261 | 1:100 |
| Lymphocyte antigen 6A-2/6E-1 | R718 | D7 | BD Biosciences | 567733 | 1:400 |
| IgD | APC-H7 | 11-26c.2a | BD Biosciences | 565348 | 1:200 |
| CD4 | APC-H7 | GK1.5 | BioLegend | 560181 | 1:400 |

**Supplementary Table 4:** List of single stains used for unmixing of fluorochromes.

| Fluorochrome | Cells/Beads* | Reference Control |
| --- | --- | --- |
| BUV395 | Beads | CD45R |
| Fixable Viability Stain 440UV | Cells | Live/Dead |
| BUV563 | Beads | CD38 |
| BUV615 | Beads | CD80 |
| BUV661 | Beads | CD326 |
| BUV737 | Beads | CD21 |
| BUV805 | Beads | CD23 |
| BV421 | Beads | CD267 |
| V450 | Beads | CD44 |
| BV480 | Cells | S6P / Streptavidin substitute |
| BV510 | Beads | IgG1 |
| BV510 | Beads | IgG2a/b |
| BV510 | Beads | IgG3 |
| BV605 | Beads | CD279 |
| BV650 | Cells | S6P / Streptavidin substitute |
| BV711 | Beads | CD183 |
| BV750 | Beads | CD138 |
| FITC | Beads | IgA |
| RB545 | Beads | CD86 |
| RB613 | Cells | RBD / Streptavidin substitute |
| RB705 | Cells | CD62L |
| RB744 | Cells | RBD / Streptavidin substitute |
| RB780 | Beads | CD19 |
| RY586 | Beads | CD93 |
| PE-CF594 | Beads | CD185 |
| NovaFluor™ Yellow 660 | Cells | IgM |
| PE-Cy5.5 | Cells | CD8a |
| PE-Cy7 | Beads | TIGIT |
| AF647 | Beads | T- and B- Cell Activation Antigen |
| APC | Beads | NK-1.1 |
| APC | Beads | TER-119 |
| APC | Beads | F4/80 |
| APC | Beads | Ly-6G/Ly-6C (Gr-1) |
| APC | Beads | CD11c |
| R718 | Cells | Lymphocyte antigen 6A-2/6E-1 |
| APC-H7 | Beads | IgD |
| APC-H7 | Beads | CD4 |

\* Single stains of each antibody were prepared using either cells or beads (splenocytes, bone marrow-derived lymphocytes, or UltraComp eBeads™). Beads were selected as single-color reference controls when they produced spectral unmixing comparable to that of cell-based controls. For antigen baits, a “dummy” control using antibodies against abundantly expressed markers conjugated to the same
fluorochrome and from the same manufacturer provided satisfactory results. Accurate spectral unmixing was confirmed with single-stained cells, and autofluorescence unmixing enhanced population separation.

**Supplementary Table 5:**
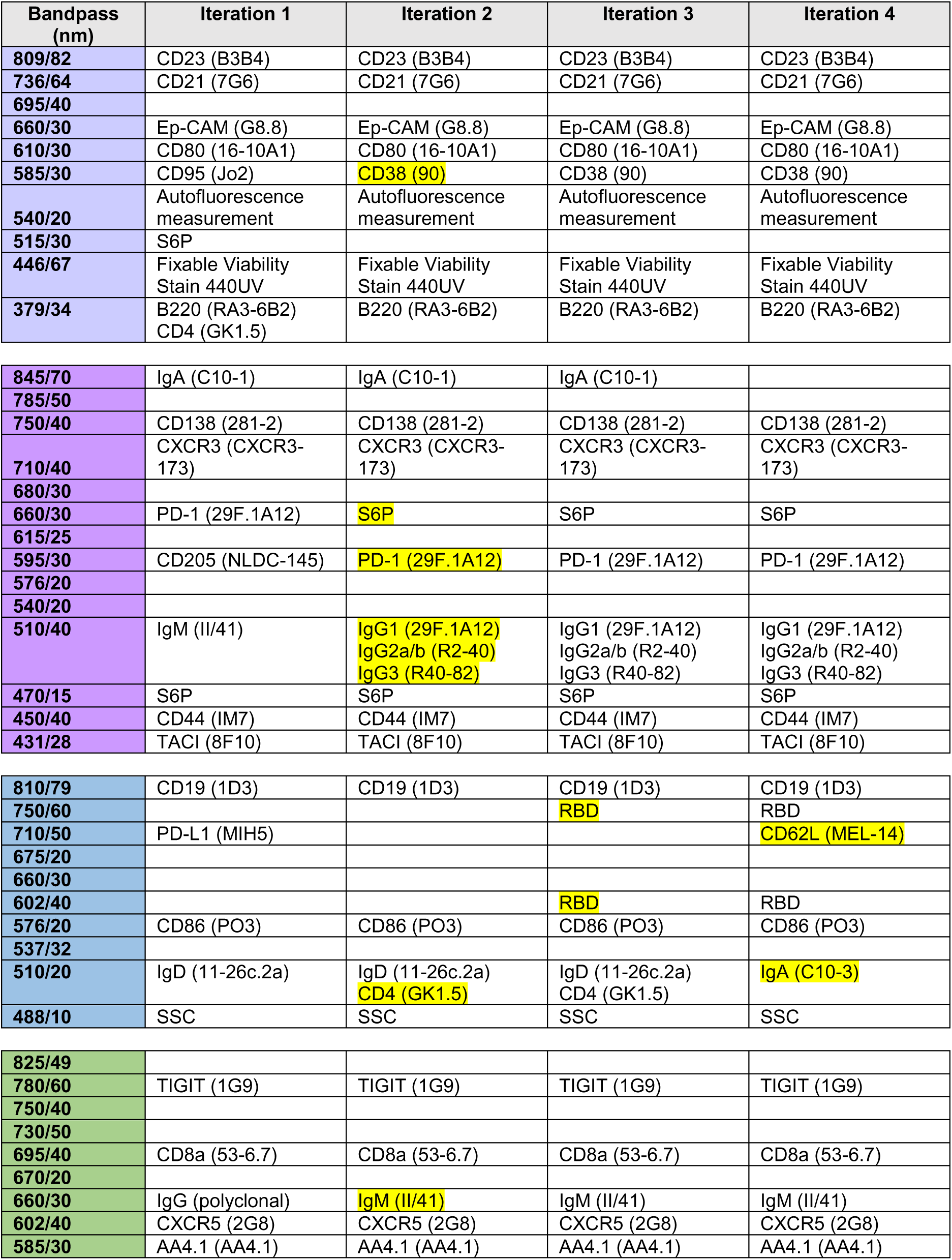
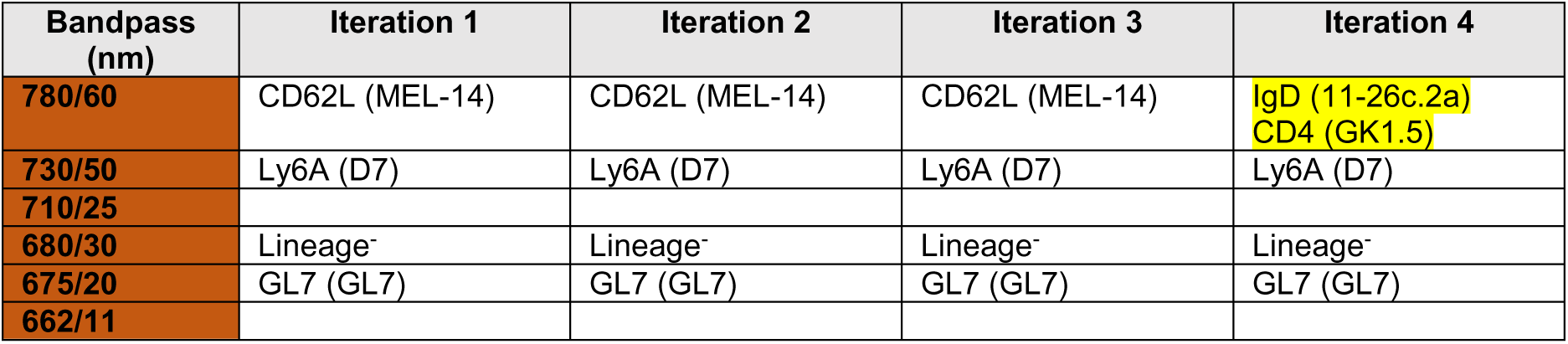
Panel design and optimization.

**Supplementary Table 6:**
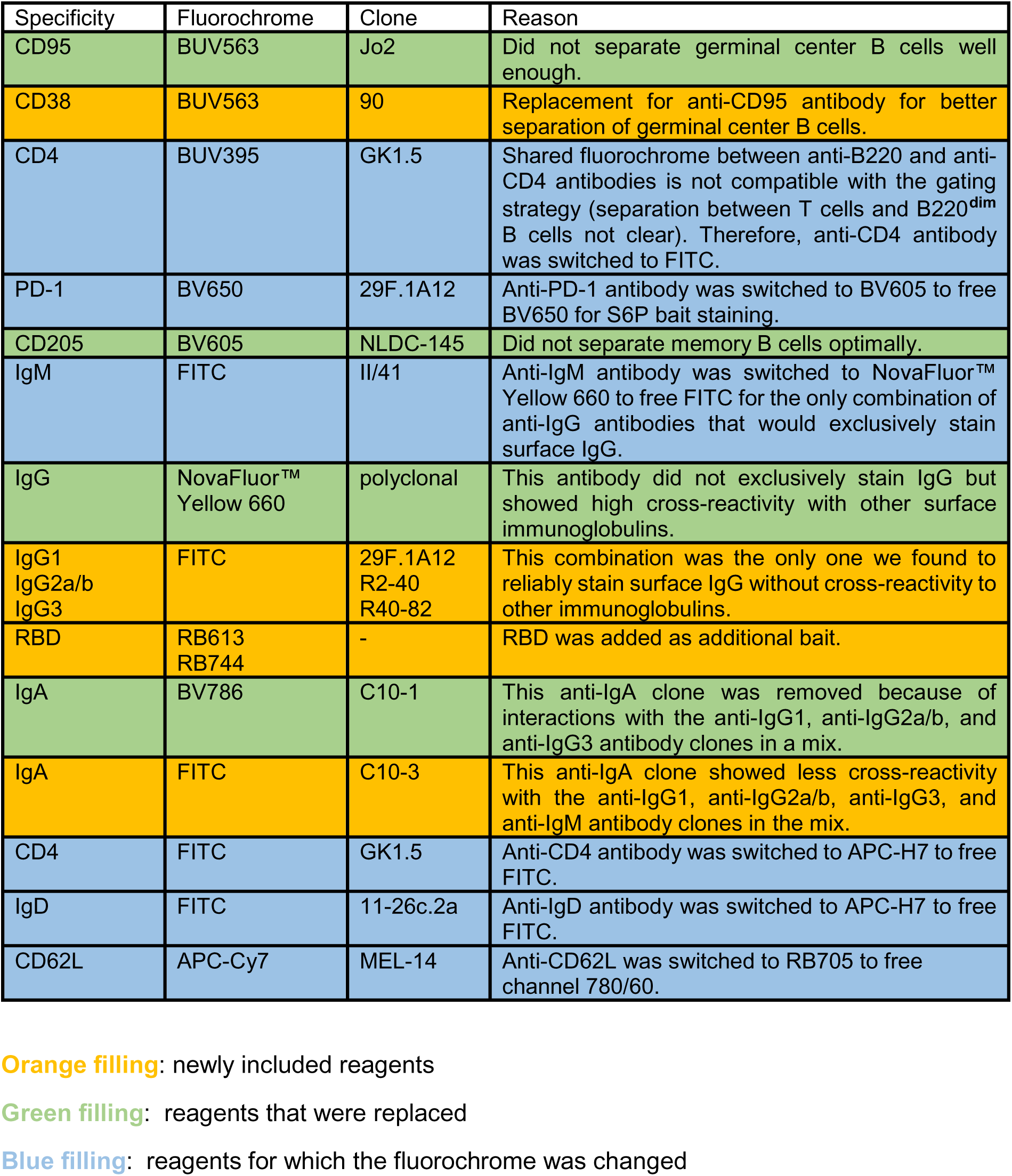
Reasons underlying panel optimization and iteration decisions.

| Specificity | Fluorochrome | Clone | Reason |
| --- | --- | --- | --- |
| CD95 | BUV563 | Jo2 | Did not separate germinal center B cells well enough. |
| CD38 | BUV563 | 90 | Replacement for anti-CD95 antibody for better separation of germinal center B cells. |
| CD4 | BUV395 | GK1.5 | Shared fluorochrome between anti-B220 and anti-CD4 antibodies is not compatible with the gating strategy (separation between T cells and B220<sup>dim</sup> B cells not clear). Therefore, anti-CD4 antibody was switched to FITC. |
| PD-1 | BV650 | 29F.1A12 | Anti-PD-1 antibody was switched to BV605 to free BV650 for S6P bait staining. |
| CD205 | BV605 | NLDC-145 | Did not separate memory B cells optimally. |
| IgM | FITC | II/41 | Anti-IgM antibody was switched to NovaFluor™ Yellow 660 to free FITC for the only combination of anti-IgG antibodies that would exclusively stain surface IgG. |
| IgG | NovaFluor™ Yellow 660 | polyclonal | This antibody did not exclusively stain IgG but showed high cross-reactivity with other surface immunoglobulins. |
| IgG1<br>IgG2a/b<br>IgG3 | FITC | 29F.1A12<br>R2-40<br>R40-82 | This combination was the only one we found to reliably stain surface IgG without cross-reactivity to other immunoglobulins. |
| RBD | RB613<br>RB744 | - | RBD was added as additional bait. |
| IgA | BV786 | C10-1 | This anti-IgA clone was removed because of interactions with the anti-IgG1, anti-IgG2a/b, and anti-IgG3 antibody clones in a mix. |
| IgA | FITC | C10-3 | This anti-IgA clone showed less cross-reactivity with the anti-IgG1, anti-IgG2a/b, anti-IgG3, and anti-IgM antibody clones in the mix. |
| CD4 | FITC | GK1.5 | Anti-CD4 antibody was switched to APC-H7 to free FITC. |
| IgD | FITC | 11-26c.2a | Anti-IgD antibody was switched to APC-H7 to free FITC. |
| CD62L | APC-Cy7 | MEL-14 | Anti-CD62L was switched to RB705 to free channel 780/60. |

**Orange filling:** newly included reagents
**Green filling:** reagents that were replaced
**Blue filling:** reagents for which the fluorochrome was changed

## Notes

### Competing Interest Statement

The authors have declared no competing interest.

